# Trans-epithelial Fluid Pumping Performance of Renal Epithelial Cells and Mechanics of Cystic Expansion

**DOI:** 10.1101/727313

**Authors:** Mohammad Ikbal Choudhury, Yizeng Li, Panagiotis Mistriotis, Eryn E. Dixon, Jing Yang, Debonil Maity, Rebecca Walker, Morgen Benson, Leigha Martin, Fatima Koroma, Feng Qian, Konstantinos Konstantopoulos, Owen M. Woodward, Sean X. Sun

## Abstract

Using a novel microfluidic platform to recapitulate fluid absorption activity of kidney cells, we report that renal epithelial cells can actively generate hydraulic pressure gradients across the epithelium. The fluidic flux declines with increasing hydraulic pressure until a stall pressure, at which the fluidic flux vanishes--in a manner similar to mechanical fluidic pumps. The developed pressure gradient translates to a force of 50-100 nanoNewtons per cell. For normal human kidney cells, the fluidic flux is from apical to basal, and the pressure is higher on the basal side. For human polycystic kidney disease (PKD) cells, the fluidic flux is reversed from basal to apical with a significantly higher stall pressure. Molecular studies and proteomic analysis reveal that renal epithelial cells are highly sensitive to hydraulic pressure gradients, developing different expression profiles and spatial arrangements of ion exchangers and the cytoskeleton in different pressure conditions. These results, together with data from osmotic and pharmacological perturbations of fluidic pumping, implicate mechanical force and hydraulic pressure as important variables during morphological changes in epithelial tubules, and provide further insights into pathophysiological mechanisms underlying the development of high luminal pressure within renal cysts.

---

Many organs are made of a series of tubules lined with epithelial cells. For the human kidney, roughly one million nephrons with 30 kilometers of epithelial tubules re-absorb 180L of water per day [1]. While the absorption activity of renal epithelial cells has been studied both in vitro and in vivo [2-4], the influence of forces and hydraulic pressures during absorption has not been examined, mainly due to difficulty in controlling these variables during experimentation. Mechanical forces are recognized as important elements during cell growth, differentiation and tissue morphogenesis [5-7]. For kidney disorders such as the polycystic kidney disease (PKD), where tubular morphology of the epithelium becomes disrupted and uncontrolled expansion of the cyst eventually results, mutations in polycystins must also alter the mechanical state of the kidney epithelium [8]. To make progress, we developed a micro-fluidic device to measure trans-epithelial fluid absorption activities of the kidney epithelium while allowing for cell imaging and simultaneous control of fluid pressure, shear stress (FSS), and media chemical composition (Fig. 1a-f and Extended Data Fig. 1a). The device measures fluidic flux across the epithelium as a function of apical and basal pressures using a microcapillary (MC) connected to port 2 or 3 (Fig. 1c). The MC measures both the trans-epithelial fluid flux and the hydraulic pressure (Fig. 1c,f). It has a volume resolution of 0.31 μL and can detect pressure changes of 10 Pa. Calibration experiments are performed to obtain capillary action contribution from the MC to the basal pressure and the static pressure profiles of each device for the shear flow condition considered (Extended Data Fig. 1b,c). Flow and pressure profiles of the entire device were also validated by simulation using FEM software (Extended Data Fig. 1d-i and SM).

**Figure 1:**
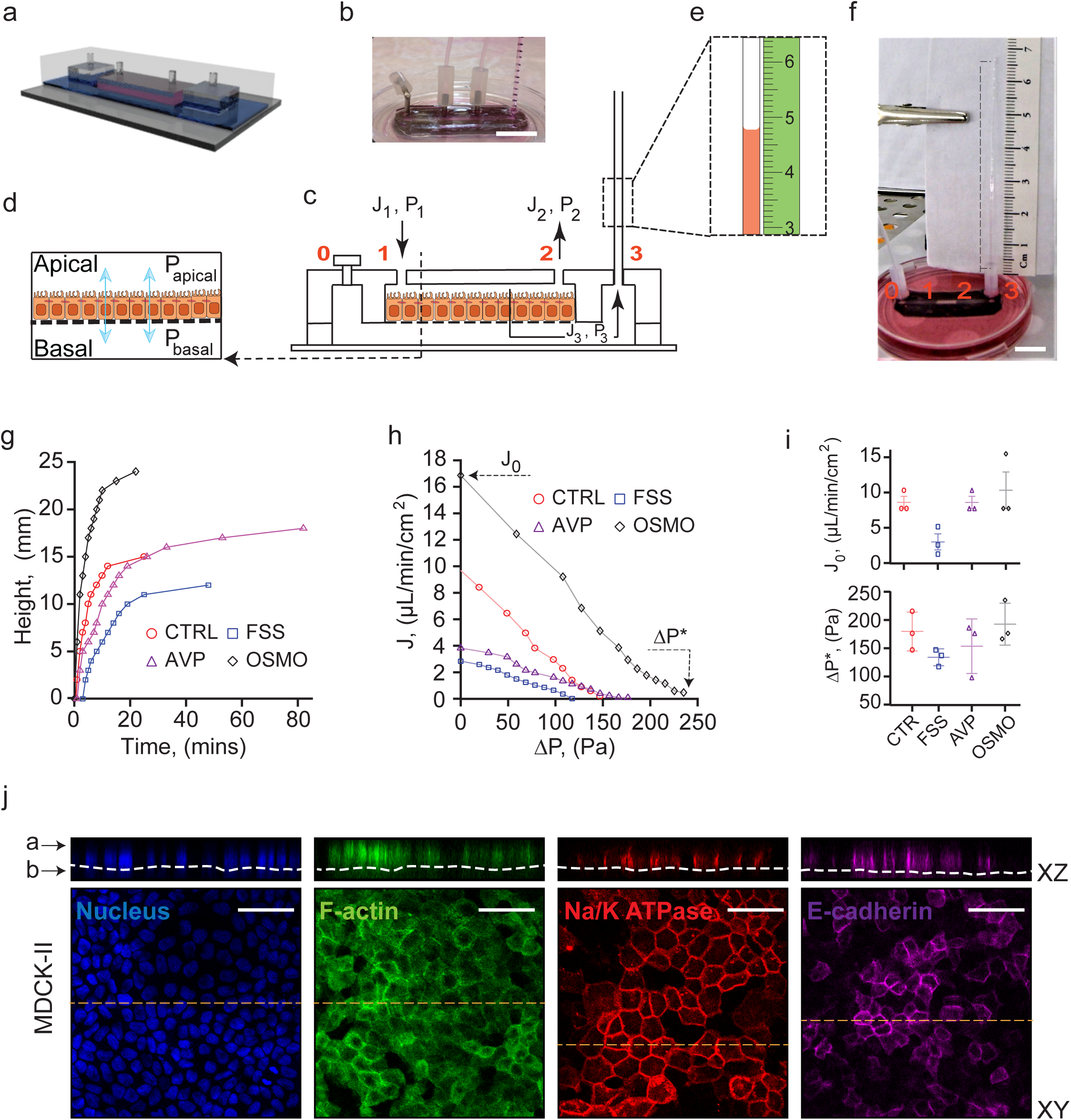
Fluid pumping performance of MDCK-II epithelium depends on mechanical, os-motic and chemical perturbation. (**a**) A schematic representation of the Micro-fluidic Kidney Pump (MFKP). (**b**) An image of the MFKP. Scale bar = 1 cm. (**c**) Longitudinal section of the device indicating four ports: 1 and 2 to access the apical channel and 0 and 3 to access the basal channel in the device. Black arrows indicate direction of fluid flux. Port 0 is closed except during seeding and cell culture. *J*_1,2,3_ are fluid fluxes and *P*_1,2,3_ are hydrostatic pressures at ports 1, 2 and 3, respectively. (**d**) Dashed arrow shows a schematic of the cross-section of the device and a polarized epithelium. *P*_apical_ and *P*_basal_ indicates hydrostatic pressures in the apical and basal channels, and arrows (blue) indicate direction of trans-epithelial fluid flow. (**e**) Dashed rectangle shows zoomed schematic of fluid flow in the microcapillary (MC) and a mm scale (green). The MC acts as a sensor to measure both *J*_3_ and *P*_3_. (**f**) A snapshot of the MFKP setup inside an incubator for time-lapsed videography. The dashed line indicates MC, and 0,1, 2 and 3 indicate ports in MFKP. Scale bar = 1 cm. (**g**) MDCK-II cells grown in MFKP generate a hydrostatic pressure gradient by pumping fluid from the apical to basal side. The height of fluid in the MC is plotted as function of time for MDCK-II epithelium in MFKP. Pumping actions of cells change under mechanical (FSS), chemical (AVP) and apical hypo-osmotic perturbations (OSMO). (**h**) Pump performance curve (PPC) of MDCK-II epithelium showing trans-epithelial fluid flux (*J*) from apical to basal versus (Δ*P* = *P*_basal_ − *P*_apical_) across the epithelium. *J*_0_ is the fluid flux at zero pressure gradient (Δ*P* = 0) and Δ*P*\* is the stall pressure when *J* = 0. Both *J*_0_ and Δ*P*\* change under FSS, AVP and OSMO perturbations. (**i**) *J*_0_ and Δ*P*\* for MDCK-II epithelium under FSS, AVP and OSMO perturbations. (**j**) Immunofluorescent (IF) images showing nucleus (blue), F-actin (green), NKA (red) and E-cadherin (purple) in MDCK-II epithelium grown in the MFKP. The dashed line (white) in the XZ panel indicate the porous membrane. The intensity projection for XZ images was recorded along the corresponding dashed lines (yellow) in the XY images. Scale bar= 25*µ*m.

When MDCK-II cells were seeded in the apical channel of the microfluidic device, cells settled on the porous membrane pre-treated with fibronectin and grew to confluence in 2-3 days. Upon further maturation, the epithelium showed classical cuboidal columnar morphology and formed a strong barrier, as tested using a dye permeation assay (see SM and Extended Data Fig. 2a-f). Visualization using immunofluorescence (IF) showed that F-actin, Na/K ATPase and E-cadherin were localized in typical fashion (Fig. 1j). Mature MDCK-II epithelium developed apical to basal fluid flow, which can be visualized as a rise in fluid height in the MC beyond the static equilibrium height (Supplementary Video 1 and Fig. 1g). The trans-epithelial fluid flux (J) from the apical to the basal channel depended on the hydrostatic pressure gradient (ΔP = P_basal_ – P_apical_) across the epithelium. J is maximal when ΔP = 0 (denoted as J_0_), and declined until a stall pressure (static head) of ΔP*∼100-250Pa was reached (Fig. 1h). This flux vs pressure curve resembled the classic pump performance curve (PPC) of mechanical fluid pumps. In contrast, for a passive filter, the pressure needs to be higher on the apical side to generate apical-basal flow, and the flux is zero when apical and basal pressures are equal. Therefore, kidney cells are active fluid pumps and our device can be considered as a microfluidic kidney pump (MFKP). The developed mechanical force is 30-100 nanoNewtons per cell, and constitutes a novel force generation mechanism that is perpendicular to the epithelium. This mechanism may be a general phenomenon for absorptive or secretory epithelia that perform fluid transport. Moreover, we found that PPC changes substantially under mechanical (FSS), chemical (addition of arginine vasopressin (AVP)) and hypo-osmotic gradient (OSMO) perturbations (Fig. 1i), indicating active regulation by cells.

To validate the trans-epithelial pressure gradient measured from our device, we examined mature polarized MDCK-II monolayer on 2D impermeable substrates (glass), which formed dynamic fluid-filled domes with elevated internal hydrostatic pressure [9]. We measured this pressure by inserting a glass micro-needle into MDCK-II domes while monitoring the curvature of an oil-media interface in the needle (Extended Data Fig. 3a-c, SM). The dome hydraulic pressure is found to be in the same range as measured in the MFKP (Extended Data Fig. 3d,e), although in domes we are unable to simultaneously monitor the trans-epithelial fluid flux. This result, together with traction force measurements of dome pressure [10], show that MDCK-II epithelium is capable of developing hydrostatic pressure of the order of 200 Pa by actively pumping fluid. To further understand the PPC curve, we also developed a mathematical model of the PPC based on active transport of an idealized solute ([11,12], see SM). If the active flux for the ideal solute depends linearly on the osmotic pressure difference across the cell apical and basal surface, then the model predicts a similar PPC that shifts with changes in apical hypo-osmotic gradient, as observed in experiments (Extended Data Fig. 6d,e).

Our device allowed us to examine molecules responsible for generating water flux. In particular, Na/K ATPase (NKA) has been implicated in directional Na^+^ transport and generation of water flow [13]. In kidney cells, NKA is polarized and accumulates in the basal-lateral surface. Blocking NKA by adding ouabain in the device’s apical channel immediately decreased trans-epithelial flux and stall pressure (Fig. 2a-c), whereas addition of Y-27367 which blocks ROCK kinase and myosin-II contractile activity, did not influence fluid flux significantly. Comparing IF images of cells in MFKP at ΔP = 0 and ΔP = ΔP* (Fig. 2d-m) showed similar F-actin distribution, but NKA showed reduced enrichment at the baso-lateral surface when ΔP = ΔP* (Fig. 2d-o). In contrast, in MDCK II domes (Supplementary Videos 3 4), NKA also showed a similar reduction in enrichment on the baso-lateral domain when the dome is stable (Extended Data Fig. 3f-q). These results indicate that cells may actively sense the apical-basal pressure difference and modulate NKA localization.

**Figure 2:**
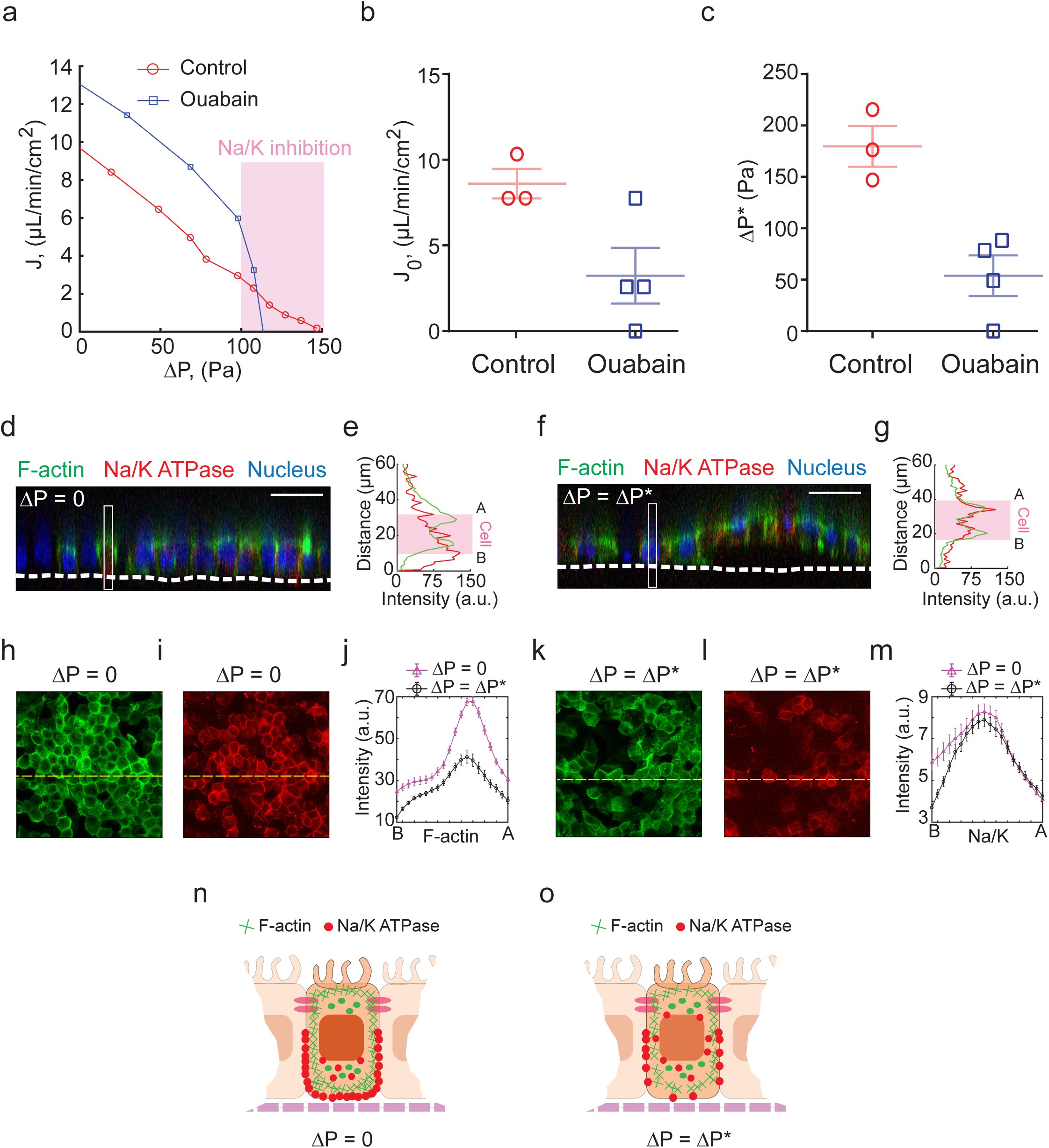
MDCK-II cells decrease the cortical expression of NKA at stall pressure. (**a**) Pump performance curves of MDCK-II cells with and without 0.2*µ*M of Ouabain. (**b-c**) Comparison of *J*_0_ and Δ*P*\* for MDCK-II epithelium in control and 0.2 *µM* Ouabain treatment. (**d**) XZ confocal section of MDCK-II epithelium in MFKP showing colocalization of F-actin (green), NKA (red) at zero hydrostatic pressure gradient (or Δ*P* = 0). The white dashed line represents the porous membrane. (**e**) IF intensity profiles of F-actin (green) and NKA (red) along the band in **d** in arbitrary units (a.u.). The pink rectangle indicates location of cell, A and B indicate apical and basal surface of the cell under consideration. (**f**) XZ confocal section of MDCK-II epithelium in MFKP showing colocalization of F-actin (green), NKA (red) in the cells at stall pressure (or Δ*P* = Δ*P*\* ≈ 200 Pa). The white dashed line represents the porous membrane. (**g**) Intensity profiles of Factin (green) and NKA (red) along the band in **f**. (**h**) Maximum intensity projected XY IF image of F-actin at Δ*P* = 0. (**i**) Maximum intensity projected XY IF image of NKA at Δ*P* = 0. The yellow dashed lines represent the line of the XZ projection in d. (**j**) Comparison of the total intensity of F-actin in five cells chosen arbitrarily along the yellow dashed line is plotted versus Z. B and A indicate the basal and apical surface of the cells. (**k**) Maximum intensity projected XY IF image of F-actin at Δ*P* = Δ*P*\*. (**l**) Maximum intensity projected XY IF image of NKA at Δ*P* = Δ*P*\*. The yellow dashed lines represent the line of the XZ projection in **f**. (**m**) Comparison of the total intensity of NKA in five cells chosen arbitrarily along the yellow dashed line plotted versus Z. (**n**) Schematic showing colocalization of F-actin (green lines) and NKA (red dots) in cells in MFKP at Δ*P* = 0. (**o**) At Δ*P* = Δ*P*\* there is a reduction in cortical NKA in the baso-lateral side.

Next, we considered whether the device is useful for understanding fluidic pumping by primary human normal kidney and ADPKD cystic cells. AQP2 (blue), Na/K ATPase (red) and F-actin (green) stains for wild type cortical cells (WTc), wild type medulla cells (WTm) and cystic cells (ADPKD) in MFKP showed the same distribution and morphology as those obtained from their corresponding immunohistochemistry images of kidney tissue sections (Extended Data Fig. 4a-d, f-k), suggesting that our device is capable of capturing the general organization of these different types of epithelia. The barrier function of the epithelia was again assessed using dye permeation assay (Extended Data Fig. 5a-e and SM).

Once grown in the device, as with MDCK-II epithelium, apical to basal fluid flux was observed as a function of hydrostatic pressure gradient in WTc and WTm epithelium, resulting in a similar PPC (Fig. 3g). J_0_ and ΔP* was higher in case of WTc as compared to WTm (Fig. 3j-o). Unlike absorptive function of normal kidney cells, ADPKD cystic cells had a secretory phenotype even though there is no dramatic difference in typical markers of apical-basal polarity [9,14]. The reversal of fluid flux in cystic cells remains unexplained and the underlying physical parameters of reversal have not been previously quantified. When the MC is connected to port 2 instead of port 3, we observed fluid rises in the MC beyond the equilibrium height, indicating basal-to-apical fluidic pumping (Supplementary Video 2). In both normal and ADPKD cells, the trans-epithelial fluid flux (J) and the PPCs are modulated by mechanical (FSS), chemical (AVP) and apical hypo-osmotic (OSMO) perturbations. These changes were quantified by plotting J_0_ and ΔP* under different conditions (Fig. 3j-o).

**Figure 3:**
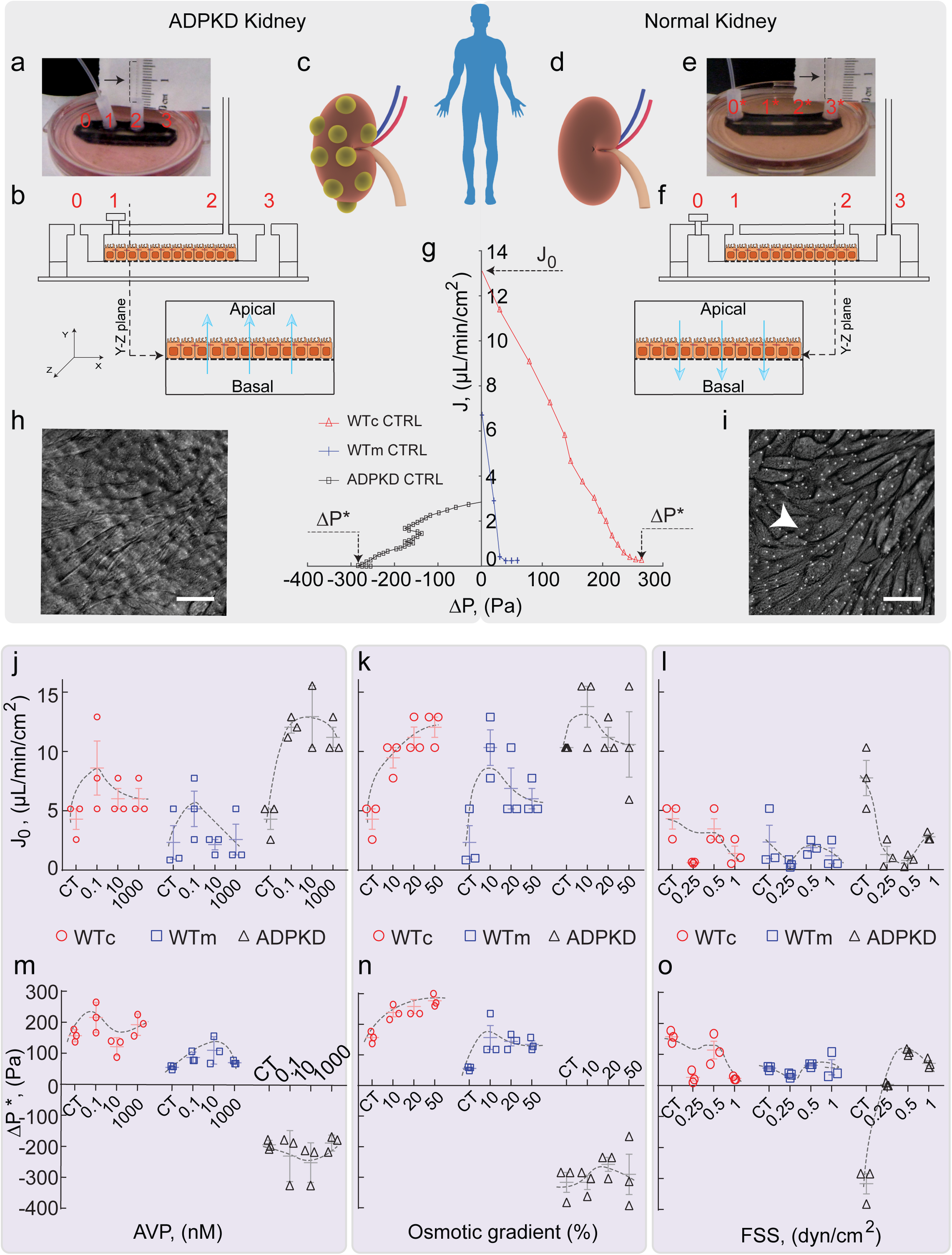
Cystic cells derived from ADPKD patients pump fluid in the opposite direction as normal human kidney cells. (**a**) A snapshot of MFKP with MC connected to port 2. Port 1 is closed. (**b**) Schematic of the setup, which allows measurement of trans-epithelial fluid flux from the basal to apical chamber. (**c**) Schematic representation of a cystic kidney derived from ADPKD patients. (**d**) Schematic representation of a kidney derived from normal human patients. (**e**) The MC is connected to port 3 in the MFKP. Port 1 is closed. (**f**) This setup enables measurement of trans-epithelial fluid flux from apical to basal chamber. (**g**) Comparison of the pump performance curves of WTc, WTm and ADPKD epithelium grown in MFKP. The trans-epithelial fluid flux (*J*) decreases with increase in hydrostatic pressure gradient (Δ*P*) across the epithelium. *J*_0_ is the fluid flux at zero pressure gradient across the epithelium (Δ*P* = 0) and Δ*P*\* is the stall pressure when *J* = 0. (**h**) DIC image of ADPKD cells forming a mature epithelium on porous membrane in the device. Arrow indicates pores (1*µ*m) in the membrane. (**i**) DIC image of WTc cells forming a mature epithelium on the porous membrane in the device. Comparison of *J*_0_ in WTc, WTm and ADPKD cells with varying AVP (**j**), hypo-osmotic gradient (**k**) and fluid shear stress (**l**). Comparison of Δ*P*\* in WTc, WTm and ADPKD cells with varying AVP (**m**), hypo-osmotic gradient (**n**) and fluid shear stress (**o**). For ADPKD cells, the fluid shear stress is applied on the basal side of the epithelium.

Our device revealed that for WTc, WTm and ADPKD cells, J_0_ and ΔP* showed variable response to basolateral treatment of AVP (Fig. 3j,m). The observed massive surge in basal-to-apical fluid pumping in ADPKD epithelium matches studies in mouse models of ADPKD, indicating cysts grow bigger with AVP treatment [15]. ΔP* however remained constant at −300 Pa for ADPKD cells. Under apical hypo-osmotic treatment, proximal tubule cells (WTc) changed their pumping performance by increasing both J_0_ and ΔP*. WTm cells also increased J_0_ and ΔP* with apical hypo-osmotic shock (Fig. 3k,n). For ADPKD cells, while J_0_ increased with decreasing basal osmolarity, ΔP* didn’t change and remained constant around −300 Pa (Fig. 3n). The converging nature of the PPC to a constant ΔP* for ADPKD epithelium (Extended Data Fig. 6c) cannot be explained by the simple active flux model (Extended Data Fig. 6f and SM), suggesting differential mechanisms of regulation during fluidic pumping in wild type and ADPKD epithelium.

In both WTc and WTm cells, apical FSS for 5 hours did not change J_0_ or ΔP* significantly (Fig. 3l,o). At 1 dyn/cm^2^, both J_0_ and ΔP decreased for both WTc and WTm as compared to control cells. However, in case of ADPKD, even though increasing FSS resulted in a decrease in the average J_0_, the fluidic flux direction reversed under FSS (Fig. 3l) with ΔP* going from −300 Pa to 100 Pa (Fig. 3o).

In ADPKD kidney, the progressive growth of fluid filled cysts leads to an increase in total kidney volume [16]. We have shown that a hydrostatic pressure gradient (ΔP) is developed during fluid pumping, and the cyst wall must sustain a pressure of ∼300 Pa. In the organ, this pressure points from the lumen towards the interstitium, and could drive cyst expansion. The regulation of ΔP* is important for understanding kidney morphogenesis. The Food and Drug Administration (FDA) approved Tolvaptan (TVP), which decreased the total kidney volume [16]. Tolvaptan is a V2R antagonist and has been shown to decrease cAMP levels [18]. We treated mature ADPKD epithelium in MFKP with 1 nM TVP on the basolateral side for 1 hour and measured PPC. Interestingly, TVP decreased both J_0_ and ΔP* of the ADPKD epithelium as compared to the control (Fig. 4a-c), confirming that the cAMP pathway can modulate fluid flux and ΔP*. To obtain a molecular profile of kidney cells during fluidic pumping, we performed qPCR measurements for aquaporins (AQP1, AQP3 and AQP5), ion-pumps and exchangers (Na/K ATPase, NHE1, NKCC1, NKCC2, CFTR), and tension sensitive Ca^2+^ channels (TRPM7 and TRPV4), which are all potentially involved in regulating water and ion transport, and mechano-sensation. Heatmaps indicating the expression of mRNAs extracted from WTc, WTm and ADPKD cells grown on permeable substrate (MFKP) and on impermeable substrate (tissue culture treated polystyrene dishes) show substantial expression differences (Extended Data Fig. 7a). Except for TRPM7 and TRPV4, expressions of all other genes were higher in cells grown in MFKP as compared to that on impermeable substrate. Moreover, using our device, we can quantify changes in genes expression as a function of ΔP. We collected cells from MFKP under two conditions- ΔP = 0 and ΔP = ΔP*. Fig. 4 (e,g,i) show that when exposed to stall pressure ΔP = ΔP*, WTm cells decreased the expressions of AQP1, AQP5, ATPA1, SLC12A1, SLC12A2, CFTR, TRPM7 and TRPV4. ADPKD cells did not respond to ΔP*, where expressions of these genes either remained constant or increased slightly. IF images of F-actin and NKA in the MFKP device corroborates the qPCR results at ΔP = 0 and ΔP = ΔP*. The total intensities of NKA in WTc, WTm and ADPKD epithelia under the two conditions were also consistent with the mRNA readings from qPCR (Fig. 4d,f,h). However, spatial arrangement of F-actin and NKA in WTc, WTm and ADPKD epithelia showed significant differences at ΔP = 0 and ΔP = ΔP* (Extended Data Fig. 8-10). For all the three human primary cell types, F-actin was generally the highest at the cell basal surface at ΔP = 0, but the F-actin stress fiber density was significantly decreased at ΔP = ΔP* (Extended Data Fig. 8-10c- f,o,q). At ΔP* cells showed increase in F-actin intensity at the cell-cell junctions in WTc and ADPKD cells but not for WTm cells. Interestingly, ΔP* disrupted the apico-basal polarization of NKA in WTm cells (Extended Data Fig. 9c-f,p,r-t). This depolarization effect was completely absent in WTc but was subtle in ADPKD cells exposed to ΔP*. This indicates that collecting duct cells (WTm population), may regulate trans-epithelial ion and fluid pumping by sensing hydrostatic pressure gradient.

**Figure 4:**
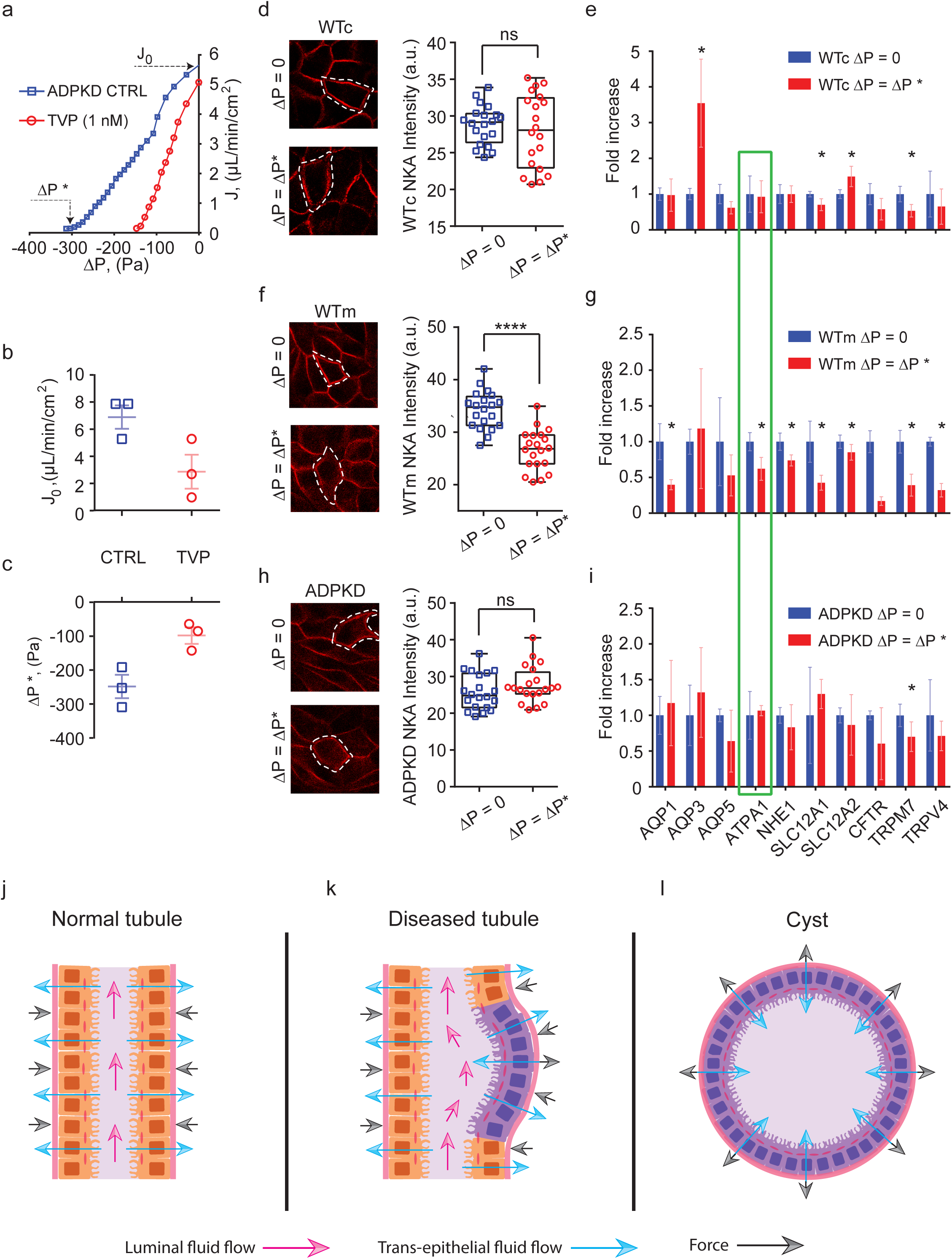
Cystic cells have defects in sensing hydrostatic pressure as compared to normal cells. (**a**) Comparison of fluid pumping performance of untreated (CTRL) ADPKD cells and ADPKD cells treated with 1 nM Tolvaptan (TVP) for 1 hour in the basolateral domain in MFKP. (**b**) Comparison of *J*_0_ in CTRL and TVP. (**c**) Comparison of stall pressure (Δ*P*\*) for CTRL and TVP. Three biological repeats are performed for each condition. (**d**) IF images of NKA in WTc cells at Δ*P* = 0 and Δ*P* = Δ*P*\* (≈ 200 Pa). Plot showing total expression of NKA in WTc cells at Δ*P* = 0 and Δ*P* = Δ*P*\* (ns = not significant, p= 0.74). Total intensity was plotted for cells picked randomly from the maximum intensity projected IF image (shown using dashed lines). (**e**) qPCR comparison of relative expression changes of genes in WTm cells at Δ*P* = 0 and Δ*P* = Δ*P*\*. (**f**) IF images of NKA in WTm cells at Δ*P* = 0 and Δ*P* = Δ*P*\*. Plot showing total expression of NKA in WTm cells at Δ*P* = 0 and Δ*P* = Δ*P*\* (p *<* 0.0001). (**g**) qPCR comparison of relative expression changes of genes in WTm cells at Δ*P* = 0 and Δ*P* = Δ*P*\*. (**h**) IF images of NKA in ADPKD cells at Δ*P* = 0 and Δ*P* = Δ*P*\*. Plot showing total expression of NKA in ADPKD cells at Δ*P* = 0 and Δ*P* = Δ*P*\* (ns, p = 0.13) (**i**) qPCR comparison of relative expression changes of gene in ADPKD cells at Δ*P* = 0 and Δ*P* = Δ*P*\*. (**j**) A schematic representation of a tubular section in the nephron of a normal human kidney. The blue arrows indicate re-absorptive fluid flux from the lumen into the interstitial space. The black arrows indicate the force due to generated pressure gradient from fluid pumping by cells. The tubular structure is stabilized by the pumping pressure difference due to apico-basal fluid flux (radially in). (**k**) During cyst initiation, cell proliferation can alter the tubule geometry and the local fluid flow pattern, results in a lowered FSS. ADPKD cells may respond by reversing the fluidic pumping direction as shown in Fig. 3o, which also reverses the pressure gradient that further destabilizes the tubule. (**l**) In mature cysts, fluid pumping elevates pressure in the cyst and aids in cyst expansion.

Our combined results offer insights into kidney fluidic pumping action and ADPKD cyst formation. Fig. 4j shows a section of a normal human kidney tubule. Blue arrows represent fluidic flux from the lumen into the interstitial space. Black arrows indicate the restoring force as a result of the fluidic pumping, arising from the apical-basal hydraulic pressure difference during pumping. Here the apical-basal pressure difference can prevent tubule expansion. During cyst initiation, due to altered morphology of the tubule, the local fluid shear stress can be lower, which leads to reversed direction of fluid pumping by ADPKD cells (Fig. 4k). This changes the direction of restoring force, which further destabilize the tubule. Together with aberrant pressure sensing response of ADPKD cells, act together to increase cyst outward pressure. These mechanical factors coupled with cell proliferation [19] can lead to gradual expansion of the cyst (Fig. 4l). Our results demonstrate that secretory and absorptive functions of epithelia can generate significant mechanical forces, and maybe a general phenomenon in tubular morphogenesis in other contexts.

