## Supplementary Information for "Trans-epithelial Fluid Pumping Performance of Renal Epithelial Cells and Mechanics of Cystic Expansion"

May 30, 2019

#### Supplemental note 1: Hydraulic pressure profile in MFKP

##### Device calibration

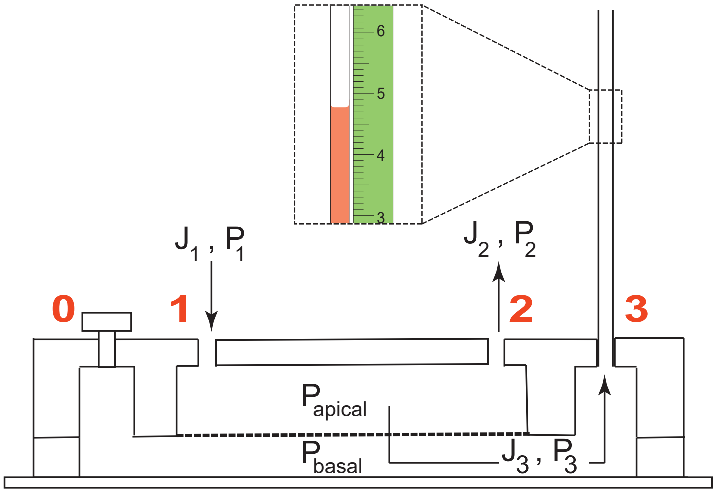


Figure 1: *Schematic of the longitudinal cross-section of MFKP without cells. Dashed box indicates a zoomed section of the fluid flow in the MC and a mm scale. $J_1$ and $P_1$ are the fluid flux and hydrostatic pressure at port 1, $J_2$ and $P_2$ are the fluid flux and hydrostatic pressure at port 2, $J_3$ and $P_3$ are the fluid flux and hydrostatic pressure at port 3.*

The hydraulic pressure profile in the apical channel of MFKP was measured under different physiologically relevant fluid shear stress (FSS) and hydrostatic pressure gradients across the device. Fluid flow rate ($Q$) in the apical channel was changed using a syringe pump connected to port 1 and FSS was calculated using:

$$\tau = \frac{6\mu Q}{ab^2} \tag{1}$$

where $\tau$ is the fluid shear stress, $\mu$ is the fluid viscosity, $Q$ is the flow rate, $a$ is the width and $b$ is the height of the channel. Fluid flow in the apical channel is due to the pressure gradient developed by the syringe pump between port 1 and 2 in order to maintain a constant flow rate (SI Figure 1). Here, $J_1$, $J_2$ and $J_3$ are the fluid fluxes through port 1, 2 and 3 and $P_1$, $P_2$ and $P_3$ are hydrostatic pressures at port 1, 2 and 3 respectively. From mass balance, the fluid fluxes through the ports are linked according to Eq. 2:

$$J_1 = J_2 + J_3 \tag{2}$$

$J_1 \equiv Q$ is a known constant that is maintained by the syringe pump. $P_1$, however is unknown. The average pressure in the apical channel ($P_{\text{apical}}$) is greater than the basal channel ($P_{\text{basal}}$), which drives the apical-to-basal fluid flux, $J_3$. When port 0 is closed, this fluid rises into the microcapillary (MC) at port 3. Here MC acts as a sensor to measure both the fluid flux $J_3$ and hydrostatic pressure $P_3$ at port 3. At steady state, $P_{\text{apical}}$ is equal to $P_{\text{basal}}$ and $J_3 = 0$. The height of fluid ($h$) in the MC at steady state was used to calculate the $P_3$ using Eq. 3, which is also equivalent to the average basal hydrostatic pressure $P_{\text{basal}}$.

$$P_3 = \rho g h \tag{3}$$

while maintaining the same FSS, $P_{\text{apical}}$ can be changed by changing the exit pressure at port 2 ($P_2$). $P_2$ was varied by changing the height of the reservoir connected to port 2. By measuring the fluid height $h$ in port 3 at $J_3 = 0$, $P_{\text{apical}}$ was measured under two different FSS and multiple ($P_2$) conditions (Extended Data Fig. 1b). Horizontal error bars indicate same-device variation (mean $\pm$ standard deviation, n = 3) and vertical error bars indicate device-device variation (mean $\pm$ standard deviation, n = 5). MFKP can therefore be used to apply a variety of physiologically relevant FSS and hydrostatic pressure to an epithelium. Due to capillary action, fluid also rises in the MC without any excess pressure. We can adjust for this extra pressure by measuring capillary height. The capillary height was measured for 5 different MCs with cell culture media. Error bars indicate standard deviation (n = 5) (Extended Data Fig. 1c).

#### Fluid flux (J) vs hydrostatic pressure gradient ($\Delta P$) plot without cells

$P_{\text{apical}}$ was also measured under no FSS or $P_2$ conditions using the same technique described in the previous section. Once $P_{\text{apical}}$ is known, the fluid flux $J_3$ was plotted as a function of the the pressure difference, $\Delta P = P_{\text{basal}} - P_{\text{apical}}$ in the absence of cells. When $P_{\text{apical}}$ is greater than $P_{\text{basal}}$, apical-to-basal (A to B) fluid flux (red) decreases to zero as the system approached equilibrium i.e. $\Delta P$ = 0, which is indicated with dashed a line in the MC schematic (left) (SI Figure 2).

When $P_{\text{basal}}$ is greater than $P_{\text{apical}}$ , fluid flux in the MC reverses and Basal-to-apical (B to A) fluid flux (blue) also decreased to zero at $\Delta P$ = 0 (SI Fig. 2). Slope of the $J_3$-$\Delta P$ plot is related to the permeability of the porous membrane. The slope for the A-to-B is higher than that of B-to-A because, in case of A-to-B the fluid flow is due to both $\Delta P$ and capillary effect of the MC.

#### FEM model

To validate the experimental measurement of $P_{\text{apical}}$ in our device, the entire microfluidic device has been modeled using the COMSOL Multiphysics software. The system was divided into three domains; apical domain (A), porous membrane and basal domain (B) (Extended Data Fig. 1d). Navier-Stokes equation (Eq. 4) was used to model the fluid domain in apical and basal channels and no slip boundary condition was enforced at the walls.

$$\rho\left(\vec{u}.\nabla\right)\vec{u} = -\nabla p + \mu\nabla^2\vec{u} + \rho\vec{g}\,, \tag{4}$$

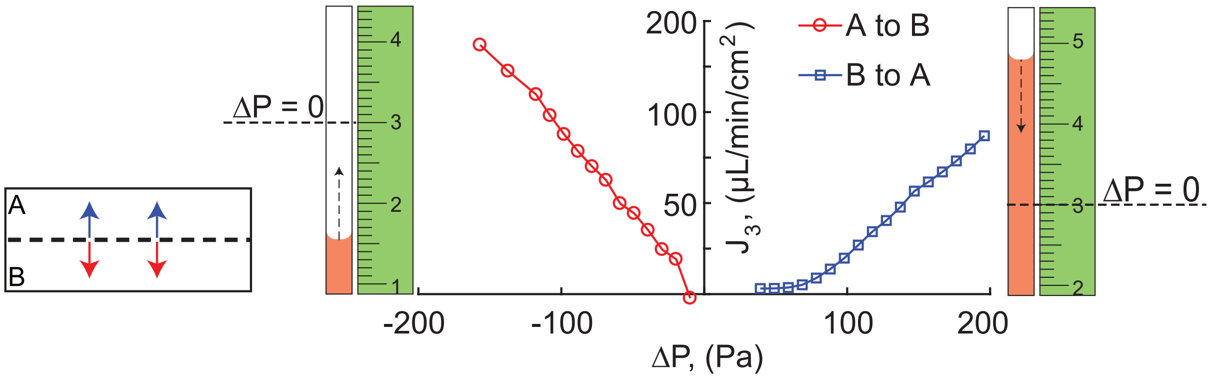


Figure 2: *Plot showing apico-basal fluid flux $J_3$ as a function of the trans-membrane hydrostatic pressure gradient $\Delta P$ without cells. The flux is in direction of the pressure gradient, and is zero when $P_{\text{basal}} = P_{\text{apical}}$.*

where $\rho$ is density of the fluid, $\vec{u}$ is the velocity vector, p is the pressure, $\mu$ is the fluid viscosity and $\vec{g}$ is the acceleration due to gravity.

The following set of boundary conditions were used:

Port 0: No Slip boundary condition as the port was blocked.

Port 1: Flux boundary condition where we prescribe flow velocity. Two flow velocities were employed, namely, 0.0047 m/s and 0.0094 m/s corresponding to fluid shear stresses 0.5 and 1 dyn/cm$^2$.

Ports 2 and 3: The height of port 2 was varied in steps of 15mm from an initial height of 10mm. This was done to recapitulate the exact experimental condition wherein a reservoir was connected to port 2. During calibrations experiments, the exit pressure $P_2$ was applied by changing the height of the reservoir.

Membrane-Channel interface: Velocity boundary conditions are used at the interface. For velocity parallel to the membrane surface, no slip zero velocity is imposed. For velocity perpendicular to the interface, a vertical velocity between 0-10$\mu$m/s can be assigned. This models the fluid flow across the porous membrane.

The computed pressure, velocity and shear stress profiles in the cross section of the device and on the surface of the porous membrane has been plotted using heat maps (Extended Data Fig. 1d-i). For both 0.5 and 1 dyn/cm$^2$, the apical channel had a longitudinal pressure gradient (along the channel direction) of about 5 Pa. Moreover, flows across the membrane did not change the apical pressure significantly ($\sim$ 3Pa). For simulations, $P_{\text{apical}}$ was calculated by taking average of the hydrostatic pressures in port 1 and 2 under all FSS and $P_2$ conditions identical to the calibration experiment (Extended Data Fig. 1b). Both experimental and simulation calibration data match closely at low $P_2$. However, at large $P_2$, the $P_{\text{apical}}$ shows a diverging trend with increase in FSS from 0.5 to 1 dyn/cm$^2$. This is because increasing the exit pressure $P_2$ causes an additional increase in the average pressure in the apical channel. Heatmaps show that the pressure $(P)$, fluid velocity $(v)$ and FSS $(\tau_{xy})$ are uniform along the XZ-crosssection (Extended Data Fig. 1e). The hydrostatic pressure on the apical channel is higher than that of the basal channel. The fluid velocity is maximum at the center of the apical channel and both velocity and FSS are uniform along the width on top the porous membrane (Extended Data Fig. 1f).

In order to validate whether $P_3$ is equal to $P_{\text{basal}}$, we plotted the pressure profile along the apical and basal channel in the YZ-axis. For both FSS 0.5 and 1 dyn/cm$^2$, hydrostatic pressure below the membrane showed minimal spatial variation, and was similar to that of port 3 (Extended Data Fig. 1g-i), indicating that Eq. (3) is accurate for measuring $P_3$. In summary, device has been calibrated experimentally and by using FEM model for hydraulic pressure profiles in both the apical and the basal channels. Once calibrated, the device accurately reports $P_{\text{apical}}$ and $P_{\text{basal}}$ within of $\pm 10$ Pa.

#### Supplemental note 2: Barrier Strength of the epithelium in MFKP

Trans-epithelial electrical resistance (TEER) is considered as the the gold standard for estimating epithelial barrier strength [3]. However, in case of MFKP, off-the-shelf electrodes do not fit in the apical and basal channel in the device. Therefore, TEER values were recorded using standard Ag/AgCl electrodes by seeding cells on membrane filters pre-treated with fibronectin. MDCK-II cells showed a sharp rise in normalized TEER value of 220 $\Omega/\text{cm}^2$ in 3-4 days post confluence. This value then plateaued to 180 $\Omega/\text{cm}^2$ with time (SI Fig. 3a). This indicates that MDCK-II cells form a strong barrier post confluence. In the MFKP the barrier strength was quantified by using a FITC-conjugated dextran dye of MW 2000 kDa. The dye was dissolved in cell culture media at $5\%$ v/v ratio and added to the apical channel through port 1 and 2 in the device and allowed to diffuse into the basal channel. Stacks of confocal fluorescence images, $20\mu m$ apart, were taken spanning the basal and apical channels. The diffusion of the dye across the epithelium to the basal channel was measured by quantifying the temporal variation of the average relative intensity of the images. Upon addition of the dye into the device without cells, the intensity quickly increased within a minute of adding the dye to the apical channel, keeping the basal channel closed. Within 5 minutes the dye diffused completely into the basal channel. The dashed line was used to indicate the position of the porous membrane that separates the basal and apical side (Extended Data Fig. 2a). This indicates that dye diffusion across the porous membrane is relatively quick ($<$10min). For MDCK-II epithelium 2 days post confluence, it took about 3 hours for the dye to diffuse into the basal side from the apical side, indicating relatively poor barrier strength (Extended Data Fig. 2b). In this case, comparison of the images of the basal and apical channel in the MFKP at $T = 0$hr and at $T = 3$hr indicated slow diffusion of the dye across the epithelium. However, For MDCK-II epithelium that is 2 weeks post confluence, the dye did not diffuse from the apical to basal channel at all. The average intensity of the images in the basal side remained the same after 3 hours post injection of dye into the apical channel (Extended Data Fig. 2c). In this case, comparison of the fluorescence images of the basal and apical channel in MFKP at T = 0 hour and at T = 3 hours indicated no diffusion of dye across the epithelium.

For human primary cells from both non-cystic and cystic kidneys, tissue samples were dissected from the cortex, medulla, or cortical cyst wall (see methods). The cells were labelled as wild type cortex (WTc), wild type medulla (WTm) and cystic (ADPKD) for all experiments. The cells were not passaged thereafter to avoid fibroblast overgrowth. The TEER values for WTc cells seeded on membrane filters reached a peak of 539 $\Omega/\text{cm}^2$ after 8 days post confluence and then plateaued (SI Fig. 3b). However, for both WTm and ADPKD cells the TEER values increased rapidly post confluence and plateaued at 180 and 350 $\Omega/\text{cm}^2$ (SI Fig. 3c,d). This indicates that both normal human kidney and PKD cystic cells exhibit strong barrier function on porous substrates. The steady state TEER value for some batch of cells reached as high as 854.22 $\Omega/\text{cm}^2$ for WTc cells (plot not shown) and 526.5 $\Omega/\text{cm}^2$ for WTm cells (two biological repeats for both WTc and WTm cells). However, sample-to-sample variability was higher for WTm cells. To measure the barrier strength of primary cells in MFKP, cells were seeded at high density by thawing directly from frozen vials (see methods). Cells grew to confluence in 2-3 days and then MFKPs were perfused with media every day

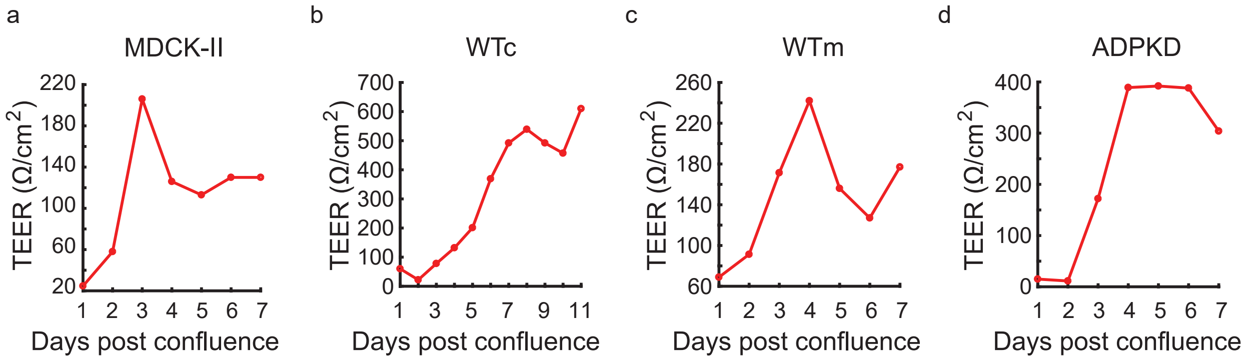


Figure 3: *Variation of the trans-epithelial electrical resistance (TEER) with the number of days post confluence for (**a**) MDCK II cells, (**b**) WTc cells, (**c**) WTm and (**d**) ADPKD cells.*

for the next 2 weeks to form a strong barrier between the apical and basal channel. In all three cell types, the FITC dye did not diffuse from the apical to basal channel. The average fluorescence intensity of the basal channel remains the same as without dye after 3 hours post injection in the apical channel (Extended Data Fig. 5a-c). The dashed white line is used to indicate the position of the porous membrane that separates the basal and apical channel. The variability of fluorescence intensity in the same device and from device to device was low (Extended Data Fig. 5g-i). Horizontal error bars indicate same-device variability and vertical error bars indicate device-to-device variability (mean $\pm$ standard deviation, n = 3). In all three types of epithelia, fluorescence images of the basal and apical channels in MFKP at $T = 0$ and at $T = 4$hr indicated no diffusion of the dye through these epithelia as shown in representative images (Extended Data Fig. 5d-f).

#### Supplemental note 3: Simple theoretical model can explain increase in $J_0$ and $\Delta P^*$ with apical hypo-osmolarity

Our experiments showed that the epithelial layer can actively pump fluid across the epithelium from the apical to basal sides. To mathematically describe this fluid pumping, we consider a steady-state description and model the flow of water across the epithelium, driven by the passage of an idealized charge-neutral solute. We assume that cells are stationary without translation or deformation. Each cell in the monolayer can be approximated as a cylinder with radius $R_0$ and height $L$ (Extended Data Fig. 6d). The cell has an apical end (a) and a basal end (b). The direction points from the basal end to the apical end of a cell is defined as the $x$-direction. $x = 0$ is the basal position while $x = L$ is the apical position. We assume that the fluid flow and solute diffusion and convection only happens in the $x$-direction within each cell. Those happen through gap junctions are neglected.

In cylindrical coordinates, the Stokes' equation for cytoplasmic water flow in the cell is

$$0 = -\frac{dp}{dx} + \mu \frac{1}{r}\frac{d}{dr}\left(r\frac{dv}{dr}\right) , \tag{5}$$

where $p$ is the intracellular hydrostatic pressure, $\mu$ is the dynamic viscosity of the fluid, and $v$ is the velocity of the flow in the positive $x$ direction. The spatial profile of $v$ is

$$v(x,r) = 2\bar{v}(x)\left[1 - \left(\frac{r}{R_0}\right)^2\right], \tag{6}$$

where

$$\bar{v}(x) = \frac{1}{\pi R_0^2}\int_0^{R_0} v(r,x)2\pi r dr \tag{7}$$

is the averaged velocity in each cross section. Eq. 6 results from the non-slip boundary condition at the cell cortex. Substituting Eq. 6 into Eq. 5 gives

$$0 = -\frac{dp}{dx} - \frac{8\mu}{R_0^2}\bar{v}\,. \tag{8}$$

The conservation of mass requires

$$\frac{dv}{dx} = 0\,, \tag{9}$$

which suggests that $\bar{v}$ must be a constant in $x$. In this case, $p$ must be linear in $x$ as seen from Eq. 8. We only need the pressure at the apical end, $p^{\mathrm{a}} = p|_{x=L}$, and the pressure at the basal end, $p^{\mathrm{b}} = p|_{x=0}$, to know the profile of the intracellular pressure $p$. The average velocity of the intracellular flow can also be solved from $p^{\mathrm{a}}$ and $p^{\mathrm{b}}$ by using Eq. 8, i.e.,

$$\bar{v} = -\frac{R_0^2}{8\mu L}(p^{\mathrm{a}} - p^{\mathrm{b}})\,, \tag{10}$$

so that $\bar{v}$ is not an unknown from modeling point of view.

Cross-membrane water fluxes occurs at the apical and the basal surfaces due to an osmotic gradient of the idealized solute (mainly $Na^+$). Our convention is that water fluxes are positive from outside to inside the cell. The continuity relation requires

$$\bar{v} = -J^{\mathrm{a}}_{\mathrm{water}} = J^{\mathrm{b}}_{\mathrm{water}}\,, \tag{11}$$

which gives two equations to solve for $p^{\mathrm{a}}$ and $p^{\mathrm{b}}$.

To find the water flux, a model for the solute concentration, $c$, is needed. The steady-state equation for solute diffusion is

$$\frac{d}{dx}\left(-D\frac{dc}{dx} + \bar{v}c\right) = 0\,. \tag{12}$$

So that the intracellular solute flux

$$J_c = -D\frac{dc}{dx} + \bar{v}c \tag{13}$$

must be a constant throughout the cell. This constant is determined by solute boundary fluxes at the apical and basal surface. We also assume that solute fluxes are positive inwards. At the cell apical and basal boundary, the solute flux is composed of a passive part, $J_{c,\mathrm{passive}}$, and an active part, $J_{c,\mathrm{active}}$ [5],

$$J_c|_{x=L} = -J^{\mathrm{a}}_c = -(J^{\mathrm{a}}_{c,\mathrm{passive}} + J^{\mathrm{a}}_{c,\mathrm{active}})\,, \tag{14}$$

$$J_c|_{x=0} = J^{\mathrm{b}}_c = J^{\mathrm{b}}_{c,\mathrm{passive}} + J^{\mathrm{b}}_{c,\mathrm{active}}\,, \tag{15}$$

where the passive flux follows the gradient of solute concentration across the cell membrane, i.e.,

$$J^i_{c,\text{passive}} = -g^i(c^i - c^i_0)\,, \quad i = \{\text{a}, \text{b}\}\,, \tag{16}$$

where $g$ is the coefficient of passive ion flux, $c^\text{a} = c|_{x=L}$, $c^\text{b} = c|_{x=0}$, and $c_0$ is the solute concentration outside of the cell. The expression for the active solute fluxes vary depending the types of solute, the cell type and potential active regulation by the cell, and is discussed below. Equations 14 and 15 serve as two boundary conditions for Eq. 12.

The water flux across the cell surface is determined by both the hydrostatic pressure gradient and solute osmotic gradient:

$$J^i_\text{water} = -\alpha^i(\Delta p^i - RT\Delta c^i)\,, \quad i = \{\text{a}, \text{b}\}\,, \tag{17}$$

where $\alpha$ is the coefficient of water permeation, which can be different at the apical and the basal ends of the cell, $R$ is the ideal gas constant, $T$ is the absolute temperature, and

$$\Delta p^i = p^i - p^i_0\,, \quad \Delta c^i = c^i - c^i_0\,, \quad i = \{\text{a}, \text{b}\}\,, \tag{18}$$

where $p_0$ is hydrostatic pressure outside of the cell.

Now we can examine different models of active solute flux across the cell surface. If we let

$$\begin{aligned} J^\text{a}_{c,\text{active}} &= \gamma^\text{a}_c RT(c^\text{a}_{in} - c^\text{a}_0 - \Delta\mu^\text{a}_c), \\ J^\text{b}_{c,\text{active}} &= \gamma^\text{b}_c RT(c^\text{b}_{in} - c^\text{b}_0 - \Delta\mu^\text{b}_c)\,, \end{aligned} \tag{19}$$

we get a PPC as shown in Extended Data Fig. 6e. (parallel lines). This models a simple solute pump where the flux is linearly proportional to the concentrate difference with a stall solute flux determined by $\Delta\mu^{\text{a,b}}_c$ [6]. This model gives an increasing fluidic pumping stall pressure as a function of decreasing osmolarity of the apical fluid, as seen for WTc epithelium in Fig. 3n.

Alternatively, if the solute pumping flux has a pressure dependence, e.g.,

$$\begin{aligned} J^\text{a}_\text{active} &= \gamma^\text{a}\frac{c^\text{b} - c^\text{a}}{c_{0,r}}k_p(p^\text{a}_0 - p_{0,r}), \\ J^\text{b}_\text{active} &= \gamma^\text{b}\frac{c^\text{b} - c^\text{a}}{c_{0,r}}k_p(p^\text{b}_0 - p_{0,r})\,, \end{aligned} \tag{20}$$

we get a PPC as shown in Extended Data Fig. 6f. (converging lines). In this case, the stall pressure, $\Delta P^*$ of the epithelium is insensitive to osmolarity of the apical fluid, as seen for ADPKD epithelium in Fig. 3n.

The model described here is phenomenological in nature, and does not include electrical charges of different species of ions. There is likely a strong coupling between different types of solutes, and a full molecular model is considerably more complex. Nevertheless, the model demonstrates that the basic physics of fluid flow across the epithelium coupled with active solute flow should produce active pumping behavior, as seen in experiments. Moreover, the observed stall pressure is probably due to a combination of active-flux dependence on osmolarity and pressure regulation of active flux. This is supported by observed changes in the localization of NaK ATPase (NKA) as a function of pressure. Therefore, a full molecular model will require understanding of how hydraulic pressure regulates active solute flux.

#### Supplemental note 4: MDCK-II Domes as three dimensional epithelial pressure vessels

##### Validation of $\Delta P$ measured in MFKP using MDCK-II domes

To validate the apical-basal pressure difference measured for MDCK-II epithelium in MFKP, we examined fluid-filled domes seen in mature polarized MDCK-II monolayer on 2D impermeable substrates (glass). Domes (or blisters) are likely developed due to trans-epithelial pumping of ions and water following a similar mechanism. The three dimensional hemi-spherical shape is sustained by the hydrostatic pressure gradient developed ($\Delta P_{\mathrm{dome}}$) [1]. We measured the $\Delta P_{\mathrm{dome}}$ by inserting a glass micro-needle-based pressure sensor into MDCK-II domes (Extended Data Fig. 3a,b). The micro-needle has an oil-water interface with a known surface tension and the other end of which is connected to a pressure sensor (Extended Data Fig. 3c). By measuring the interfacial curvature, the hydrostatic pressure in the dome can be calculated using Young-Laplace equation (Eq. 21). Further details of the exact working principle of this device is described elsewhere [2].

$$P_1 - P_2 = \frac{2\gamma}{R}\,, \tag{21}$$

where $P_1 - P_2$ is the pressure difference across the oil-media interface, $\gamma$ is the oil-media surface tension for the interface, and $R$ is the mean radius of curvature of the oil-media interface (Extended Data Fig. 3c). Since $P_1$ is the pressure inside the dome, knowing the hydrostatic pressure in the media then gives $\Delta P_{\mathrm{dome}}$. $\Delta P_{\mathrm{dome}}$ in MDCK-II domes of varying size was found to be in similar range as that measured in case of the MFKP (Extended Data Fig. 3d, e). Note this method does not simultaneously measure fluid flux across the epithelium, therefore we do not know which part of the PPC curve the measurement is probing. However, the method provides estimates on upper bounds in dome pressure. These results, together with traction force measurements of pressure inside the same type of domes [4], show that MDCK-II epithelium can develop hydrostatic pressure of the order of 200 Pa.

##### Role of hydrostatic pressure gradient on the baso-lateral localization of NKA

Mature MDCK-II domes in epithelia on 2D impermeable substrates (glass) were also used to investigate the effect of hydrostatic pressure gradient on the localization of F-actin and NKA in the cells forming domes. We compared cells that have just formed a lumen, referred to here as a pre-dome or unstable ($\Delta P_{\mathrm{dome}} \approx 0$), with the cells experiencing high pressure (near stall pressure) in mature domes ($\Delta P_{\mathrm{dome}} \approx \Delta P^{*}$) (Extended Data Fig. 3f,g). Time lapse videos show that MDCK-II domes were unstable when the monolayer was immature, and were prone to collapse, likely due excessive fluid leakage (SI Video 3). However, as the monolayer matures, the domes became stable and eventually reaching a steady size due to a stall pressure (SI Video 4). We hypothesized that hydrostatic pressure gradient can change the baso-lateral polarization of NKA, which then leads to the decrease in trans-epithelial fluid flux. In both cases the epithelium was a free-standing monolayer, hence the influence of substrate focal adhesions on the localization of NKA can be ignored. Confocal reconstruction of MDCK-II pre-dome showed enrichment of F-actin in the cortex and NKA primarily on the baso-lateral domain (Extended Data Fig. 3h-k). However, in case of cells in mature domes experiencing high pressure near stall, NKA expression was depleted in the basal domain (Extended Data Fig. 3l-o). F-actin was still cortical in nature. By plotting the fluorescence intensity of each slice in the confocal stack versus the distance, we found that the overall intensity of both F-actin and NKA in the cells also decreased when the cells are experiencing near stall pressure (Extended Data Fig. 3p,q). Lattore et al. previously reported mechanical strain heterogeneity in MDCK cells forming domes of controlled sizes and cortical dilution of F-actin in super-stretched cells [4]. Therefore, to decouple the role of stretch from hydrostatic pressure, we chose to investigate relatively less stretched cells only. We found out that both stretched and unstretched cells in mature domes exhibit depolarization of NKA (SI Fig. 4). Therefore these results are further evidence that the hydrostatic pressure gradient plays an important role in NKA polarization in MDCK-II cells.

#### Supplemental note 5: Phenotypic similarity of cells in MFKP with tissue section under normal and diseased condition

Tissue sections from normal human kidney and cystic human ADPKD kidneys were compared with epithelial monolayers (wild type and ADPKD) grown in MFKP. Immunohistochemical analysis of tissue sections reveal that NKA (red) is expressed on the basolateral side of cells in both normal renal tubules and ADPKD cysts (Extended Data Fig. 4a, b). AQP2 (blue) is however enriched on the apical or sometimes sub-apical domains for both normal human kidney and ADPKD cysts (Extended Data Fig. 4c, d). The asterisk indicates the lumen side in cystic tissue section. This indicates that the usual markers of apico-basal polarity are similar in both normal tubular and cystic cells. The normal cells demonstrate regular cuboidal columnar morphology. However, cystic cells have an irregular, stretchy shape and decreased cell height (Extended Data Fig. 4a, c).

The progressive growth of fluid filled cysts leads to increase in total kidney volume, which impairs normal function of the kidney in ADPKD patients [7]. Fluid accumulates into the cyst lumen over long period of time and the hydrostatic pressure is sustained by the epithelium lining the cyst. We analyzed the fluid collected from the same cysts from which the cells for PPC experiments were extracted (Extended Data Fig. 4e). The average osmolality of the fluid collected from all the 16 cysts was 315.83 mmol/kg, and the concentrations of $Na^+$, $K^+$ and $Cl^-$ were found to be 137.19, 4.93 and 100.75 mmol/L, which is in the similar range of normal interstitial fluid [8]. Therefore, even if the cells pump ions and hence water from basal to apical side, the steady state osmolality and ion concentration were similar to interstitial fluid due to water flux into the lumen.

Immunocytochemical analysis of wild type cortex (WTc) grown in MFKP demonstrates co-localization of AQP2 (blue), NaKATPase (red) and F-actin (green). XY image represents top view of the cells on the porous membrane and XZ images show cross-sectional view of the cells along a line of interest (Extended Data Fig. 4f). The dashed white line in XZ image represents the porous membrane. Quantification of apico-basal intensity for AQP2, NKA and F-actin in WTc epithelium showed enrichment of NKA primarily in the basolateral domain of the cells indicated by white arrow on the corresponding XZ image in Fig. 3C. AQP2 intensity was low indicating absence of the protein in WTc cells, as the cortical tissue primarily is comprised of proximal tubule cells which lack AQP2 (Extended Data Fig. 4f). F-actin was fibrous on the basal side with strong enrichment in the cell-cell junctions (indicated by arrow in XZ) was observed. F-actin XZ images demonstrate that WTc epithelium exhibit the regular columnar cuboidal epithelium like that of cells in the non-cystic tissue sections (Extended Data Fig. 4a, f). In case of WTm cells, AQP2 was highly enriched in the apical domain and was also present all over the cytoplasm. NKA intensity was low and is primarily located in the basal side of the cells. F-actin was fibrous throughout the cytoplasm and strong enrichment in the cell-cell junctions was observed (marked by arrow in the F-actin XZ image (Extended

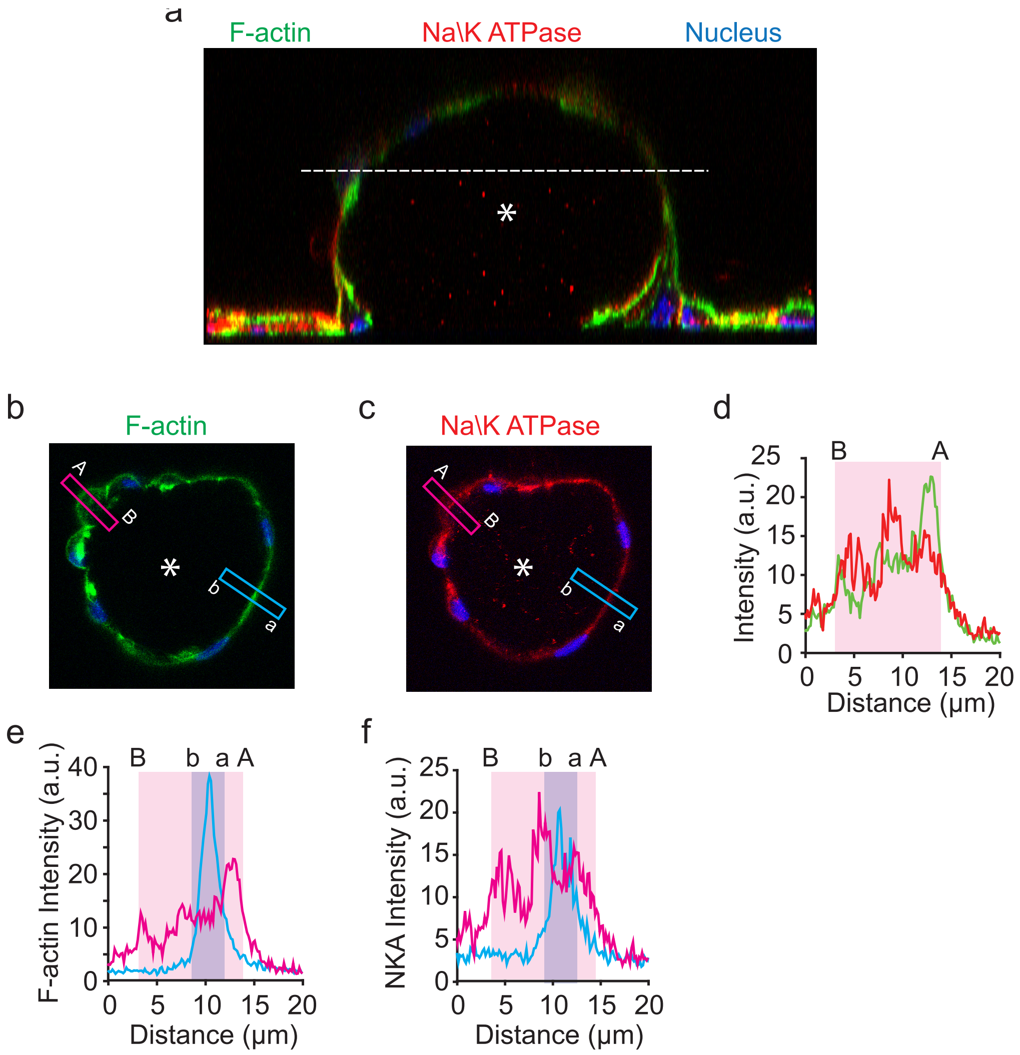


Figure 4: *(**a**) 3D confocal reconstruction of a mature MDCK-II dome stained for F-actin (green), NKA ($\alpha$ sub-unit) (red) and DNA (Blue). The asterisk indicates lumen in the dome. (**b**) Cross-sectional IF image of F-actin along the dashed line in **a**. (**c**) Cross-sectional IF image of NKA along the dashed line in **a**. (**d**) Comparison of intensity profiles of F-actin and NKA from basal to apical side of a cell in **b** and **c**. The apical and basal domains of the cell are indicated by A and B. Green line indicates F-actin intensity along the pink band in **b** and red line indicates NKA intensity along the blue band in **c**. (**e**) comparison of basal to apical intensity of F-actin in a stretched and unstretched cell in the dome in **b**. The blue line indicates the F-actin intensity along the blue band in **b**. The pink line indicates the F-actin intensity along the pink band in **b**. (**f**) Comparison of basal to apical intensity of NKA in a stretched and unstretched cell in the dome in **b**. The blue line indicates the NKA intensity along the blue band in **b**. The pink line indicates the NKA intensity along the pink band in **b**. The apical and basal domains of the unstretched cell are indicated by A and B. The apical and basal domains of the stretched cell are indicated by **a** and **b**.*

Data Fig. 4h,i)). ADPKD cells were found to have AQP2 in on the apical side in a sporadic fashion in the epithelium. NKA was enriched in the basolateral domain and is not uniformly distributed all over the epithelium indicating cell type heterogeneity. F-actin was highly fibrous in the baso-lateral domain and unlike normal cells localization in the cell-cell junction was low (Extended Data Fig. 4j,k).

#### Supplemental note 6: Normal and diseased cells on permeable and impermeable substrate

In order to investigate the influence of $\Delta P$ on the mRNA expression of the cells, we performed qPCR on ten important genes involved in regulating ion/water transport and mechano-sensation. Heatmaps indicating the expression of mRNAs extracted from WTc, WTm and ADPKD cells grown on permeable substrate (MFKP) and on impermeable substrate (tissue culture treated polystyrene dishes). The rows are normalized such that the relative concentration of across the cells lines has been shown using color code (Extended Data Fig. 7a). The genes under investigation included Aquaporins (AQP1, AQP3 and AQP5) and ion-pumps (NKA, NHE1, NKCC1, NKCC2) and ion-channels (CFTR, TRPM7 and TRPV4). Except for TRPM7 and TRPV4, the expression for all other genes was higher in cells grown in MFKP as compared to that on impermeable substrate, suggesting that the monolayer becomes much more physiologically relevant when grown in MFKP. Two biological repeats were done for each condition.

Cells plated on the impermeable substrate seemed to express a higher abundance of polycysin protein than cells plated on permeable substrate, when analyzed by western blot. Since equal loading of protein was ensured by use of a loading control ($\beta$- actin), we can be sure that an equal number of cells is represented in each lane. This leads us to believe that the cells express protein differently on the different substrates. This phenomenon may be explained by the ability of the cells to form polarized layers on the permeable membrane. The cells may be able to continue dividing for longer and therefore require more time to fully express the Polycystins. However, the cells grown on impermeable substrate do not polarize but do become confluent quickly, therefore expression of Polycystins may be higher in these cells. Polycystin levels are generally low in adult kidney cells yet it is possible to detect a weak band in each of the samples by western blot. Both Polycystin 1 and Polycystin 2 appear to be expressed in the ADPKD derived cells. This indicates that the mutation in the ADPKD kidney is not a full deletion of neither PKD1 nor PKD2 but rather appears to affect function. This phenomenon has been noted in animal models where dosage of Polycystins affects cyst growth [9] and loss-of-fucntion of Polycystin 1 is linked to cystic disease [10]. Extended data Fig. 7b-f confirms that we are able to detect Polycystin expression levels in all samples.

#### Supplemental note 7: Role of hydrostatic pressure gradient in localization of NKA in normal and diseased human primary cells.

NKA (NKA) has been implicated as the driving force behind trans-epithelial $Na^+$ transport and vectorial fluid movement in both normal and diseased cells [11]. Upon Ouabain induced inhibition of NKA in WTc cells, the equilibrium fluid flux ($J_0$) decreased from 4.3 to 1.7 $\mu$L/min/cm$^2$ and stall pressure ($\Delta P^*$) decreased from 157 Pa to 34 Pa (Extended data Fig. 8a, b). For WTm cells, overnight incubation with 274 $\mu M$ Ouabain caused a decrease of $J_0$ from 4.3 to 1.0 $\mu$L/min/cm$^2$ and $\Delta P^*$ from 56 Pa to 42 ($\Delta P^*$) (Extended data Fig. 9a, b). Ouabain caused a similar decrease in both $J_0$ and $\Delta P^*$ for ADPKD cells as well (Extended data Fig. 10a, b). Even though the direction of fluid flow in normal human renal tubular and ADPKD cystic cells are different, $J$ decreased with $\Delta P$ for all the three types of primary cells. To investigate the role of $\Delta P$ on the localization of NKA, a hydrostatic pressure difference of 200 Pa was applied to the basal channel for WT cells and to the apical channel for ADPKD cells. These cells were then fixed and stained to examine for cytoskeletal organization and NKA localization.

In case of WTc cells in control conditions (indicated by $\Delta P = 0$), F-actin is cortical in nature and primarily localized in the basal side, forming thick stress fibers (Extended data Fig. 8c,d,g). NKA is enriched in the canonical baso-lateral domain with strong enrichment on lateral domain (Extended data Fig. 8c,d,h). However, upon application of basal stall pressure (indicated by $\Delta P = \Delta P^*$), the basal stress fibers disappeared. The F-actin intensity in the cell-cell junctions was thicker in case of stall pressure as compared to control cells, indicating F-actin reinforcement in the junctions with perturbation (Extended data Fig. 8k,m,o,q). The baso-lateral polarization and the enrichment of NKA on the lateral side was same under both control and basal $\Delta P^*$ (Extended data Fig. 8k-r). The total NKA intensity was calculated by randomly choosing 20 cells in the maximum intensity projected view of both control and $\Delta P^*$ cells (Extended Data Fig. 8h,j). In case of cells exposed to basal $\Delta P^*$, The total intensity had a wide distribution as compared to the control cells (Fig. 4d). This could be the reason why the NKA intensity of individual slices decreased a bit in the lateral domain (Extended Data Fig. 8p). However the difference in the average NKA intensity under both conditions was not significant ($p = 0.74$) This is consistent with the mRNA levels of ATPA1 under both conditions (Fig. 4e).

WTm cells had thick basal F-actin stress fibers at $\Delta P = 0$, which is indicative of strong focal adhesion formation. NKA was polarized on the baso-lateral side as expected (Extended data Fig. 9c,d,g,h ). Stall pressure (indicated by $\Delta P = \Delta P^*$) on the basal side caused a loss of basal F-actin fibers but the basal-to-apical distribution was still maintained. However, $\Delta P^*$ induced a dramatic depolarization of NKA as the localization changed from baso-lateral to mostly cytoplasmic (Extended data Fig. 9e,f,i,j). This effect becomes more clear when we plotted the total intensity of each slice in the confocal stack against the distance (Extended Data Fig. 9o,p). Even though the lateral domain has more NKA as compared to the basal side in case of $\Delta P^*$ but the intensity is lower than that of control cells. As compared to the control, $\Delta P^*$ also diluted the enrichment of NKA on the lateral side (Extended data Fig. 9l,n,r). The F-actin expression in the cell-cell junctions did not change (Extended data Fig. 9k,m,q). The loss of F-actin on the basal side in WTm cells due to $\Delta P^*$ was similar to that of WTc cells but the baso-lateral dilution and depolarization of NKA is more dramatic in WTm cells (Extended data Fig. 8s,t and 9s,t). We compared the total NKA intensity of 20 cells chosen randomly in the maximum intensity projected view of both control and $\Delta P^*$ cells (Extended Data Fig. 9h,j). We found that the total NKA expression decreased with basal $\Delta P^*$ ($p < 0.0001$) (Fig. 4f). Therefore, for WTm cells, basal $\Delta P^*$ not only changed the localization of NKA but also decreased the overall expression in the cells, which is consistent with the decrease in mRNA expression of ATPA1 under similar perturbation (Fig. 4g).

Like WTc and WTm cells, cystic cells from ADPKD patients also have thick F-actin stress fibers on the basal side. The intensity of F-actin is more on the basal side as compared to the apical side (Extended data Fig. 8c,d,g, 9c,d,g and 10c,d,g). NKA is also polarized in the baso-lateral domain but the total expression is lower than that of WTc and WTm cells (Extended Data Fig. 10c,d,h and Fig. 4h). Apical $\Delta P^*$ changed the F-actin organization but not the NKA expression or localization. Unlike WTc and WTm where basal $\Delta P^*$ caused a dramatic decrease in basal F-actin stress fibers, in case of ADPKD cells, Apical $\Delta P^*$ induced a small decrease of basal F-actin stress fibers (Extended data Fig. 8g,i,o, 9g,i,o and 10g,i,o). Rather the enrichment of F-actin on the sub-apical domain was observed. Clearly, all three cell types changed their cytoskeletal organization of F-actin when exposed to a hydrostatic pressure difference. NKA localization remained baso-lateral under both conditions (Extended data Fig. 10c-f,h,j). The total intensity was also same in both conditions ($p = 0.13$) (Fig. 4h). The F-actin enrichment in the cell-cell junction and the NKA enrichment on the lateral side of the cells was also same under both conditions (Extended data Fig. 10k-n,q,r) .

### Methods:

#### Fabrication of Micro-Fluidic Kidney Pump (MFKP)

The MFKP device has four primary components- micropatterned PDMS block (Ellsworth Adhesives 4019862), porous membrane (Sigma Z681822-3EA), micropatterned intermediate layer (Ted Pella 16081-2) and glass slide (WPI FD5040-100) as shown in Extended Data Fig. 1a. The PDMS block was fabricated by mixing PDMS base and curing agent in a 10:1 ratio and pouring over an aluminum master mold, manufactured using standard micromachining. Then it was cured for 12 hours at room temperature and for 6 hours at 800 C. The central pattern has dimensions of 6mm x 2 mm x 250 $\mu m.$ Two more patterns in the PDMS block have dimensions of 2 mm x 2mm x 250 $\mu m.$ Four holes were punched in the PDMS block to create ports for media inlet and exit. The porous membrane has a thickness of 10 $\mu m$ and pore size of 1 $\mu m$, was cut into the right size before bonding it with the PDMS block. A sharp scalpel was used to make patterns with dimensions of 30 mm x 2 mm x 200 $\mu m$ in the intermediate layer. The PDMS block, the porous membrane and the intermediate layer were bonded using an RTV sealant (Sky Geek 4102963) by manually aligning them to form an assembly. This assembly was then connected to the glass slide after oxygen plasma treating (Plasma Etch PE-25) all components to make final assembly. The interfaces of and the glass slide were sealed with an RTV sealant.

#### Calibration for pressure and Flux

A syringe pump (New Era Pump Sys NE-1000) with a 30 ml syringe (Fisher Scientific 302832) filled with cell culture media was connected to ports 1 and 2 of the device to apply a range of shear stress and hydrostatic pressure in the chamber A, whereas the chamber B was filled with the same media. Port 0 was closed and port 3 was connected to microcapillary (Sigma P2049-1PAK) (Fig. 1 C, E). A mm scale paper ruler (Wintape PT-025) was glued to a holder next to the microcapillary to image the height fluid in the microcapillary using videography. As shown in Fig. 1 F, due to the pressure gradient media will flow from channel A to B, though the porous membrane and start filling the microcapillary. As the fluid rises in the microcapillary, the final height (h) was recorded using a camera (GoPro Hero7) by making time-lapsed videography of the entire setup in the incubator. Pressure in channel B can be measured by multiplying density ($\rho$) of the media and acceleration due to gravity (g) to the measured height (h) of the fluid column in the microcapillary. This is also the average pressure ($P_{\text{apical}}$) in channel A due to steady state pressure equilibration (see SM). The average pressure ($P_{\text{apical}}$) in channel A was mapped for all shear stresses and pressures $P_2$ in port 2 of the devices (Extended Data Fig. 1b). The variability in a single device was calculated by taking 3 pressure measurements in the same device and device-to-device variability was obtained by taking measurements from 5 different devices.

#### Extraction of cells from Normal and ADPKD patients

Human ADPKD cystic renal tissue from nephrectomized patients was received from the Cell Culture and Engineering Core of the Baltimore Polycystic Kidney Disease Research and Clinical Core Center. Normal human kidney samples were received from the Living Legacy Foundation and consisted of kidneys not suitable for transplant. These kidneys were disease free and within normal ranges for function (adjusted for age). For primary cell culture from both non-cystic and cystic kidneys, tissue samples were dissected from the cortex, medulla, or cortical cyst wall. The tissue sample was further reduced in size with micro-scissors in a 35 mm culture dish and washed with 0.2% collagenase A (Sigma C2139) in renal epithelial cell (REC) media (devised by Dr. Erik Schweibert of the Baltimore Polycystic Kidney Disease Research and Clinical Core Center). REC media consists of a 1:1 mixture of RenaLife Complete Medium (Lifeline Cell Technology LL-0025) and Advanced MEM medium (Fisher Scientific 12492) with 5% FBS (ThermoFisher 26140-079), 2.2% Pen/Strep (Fisher Scientific 30-002-Cl), 0.6% L-alanyl-Glutamine (Gemini Bio-products 400-106), and 0.03% Gentamicin (Quality Biological 120-098-661).The minced tissue digest was then transferred to a 15 mL falcon tube coated with 2.64% bovine serum albumin (Sigma A9574) in 1X PBS (Gibco 10010-023). Final volume of collagenase solution and tissue was 4 mL. The tubes were then incubated in a $37^0$C water bath for three hours, shaking at 125 rpm. During the incubation, tissue samples were mechanically disrupted every hour with a 2.64% BSA coated 1 mL pipette tip, by pipetting up and down 15-25 times. Following the incubation, 10 mL of cold REC media was used to inactivate enzyme reaction. Samples were centrifuged for 10 minutes at 1200 rpm, at $4^0$C, then the media aspirated, and cell pellet washed with another 5 mL of REC media, and centrifugation repeated. The cell pellet was then resuspended in 4 mL of Baltimore PKD media. Finally, 2 mL of cell suspension was plated in a T75 cell culture flask (Denville T1225) with 10 mL of REC media. Cells were expanded until confluent, then either frozen for storage or replated into the microfluidic device.

#### Culturing primary cells in the microfluidic device

Prior to seeding cells, all the devices were oxygen plasma treated to increase hydrophilicity and maintain sterility. The devices were then rinsed in 70% ethanol for 10 mins and then washed thoroughly with cell culture media and kept filled with the same media for 12 hours in an incubator maintained at $37^0$C and 5% $CO_2$. To improve cell adhesion, the channels were filled with 50 $\mu g/ml$ of fibronectin () solution for overnight at $4^0$C. The fibronectin solution was then washed off and the devices were thoroughly rinsed with cell culture media and incubated for 12 hours, before seeding cells in the chamber A of the device. MDCK-II cells were passaged at least once in growth media comprising of DMEM1x (Corning 10-013-CV), supplemented with 10% Fetal Bovine Serum (FBS) (ATCC 30-2020) and 1% Penicillin-streptomycin (ThermoFisher 15140163), before seeding in the device. However normal human wild type (WT) cells and cystic cells from ADPKD patients were not passaged in order to avoid fibroblast overgrowth but seeded directly into the microfluidic device after thawing the vials containing frozen cells. After thawing with warmed up REC media, the cell solution was gently centrifuged () at 1000 rpm for 2 mins. The cell pellet was then collected by discarding the supernatant and then resuspended in the warmed up in 200 $\mu L$ of REC media. The cells were seeded at very high density using a 1 ml syringe (BD 309628) with 18-gauge blunt tips (McMaster 75165A676) and grew to confluency over 2-3 days which was evaluated using a phase-contrast microscope. At day 1 post-seeding, the non-adherent cells were washed out by gently adding media through port 1 in chamber A. The media was exchanged every 2 days for 2 weeks post confluence to form a tight mature epithelium before preparing for experiments. The devices with incomplete monolayer as observed by phase-contrast microscope were discarded.

#### Epithelial barrier strength

The trans-epithelial electric resistance was measured using off-the-shelf TEER measurement device (EVOM-2, WPI Inc, Sarasota, FL) using Ag/AgCl chopstick electrodes (STX-2) with cells seeded on track-etched

PET membranes with 1 $\mu m$ pore diameter (Corning 353104). For each measurement, the plates were taken out of the incubator and equilibrated to room temperature for about 10 minutes in the biosafety cabinet (during this time the chopstick electrodes were equilibrated in room temperature culture media). Three measurements for each well were taken, measuring the blank transwell in between each round. The order of the wells was random each time. The reported TEER values were calculated by subtracting the average blank reading (no cells) for each set.

To test the permeability of the epithelium 5% v/v FITC-conjugated dextran (FITC-Dex, Sigma) solution with a molecular weight of 2000 kDa was added to the apical side (chamber A) of the device and the basal side sealed off. Confocal imaging (Zeiss) was used to take fluorescence images of 20 field of views from the basal to the apical side each 20 $\mu m$ apart. Stacks of images were taken every hour for 3-4 hours at the same location. The final intensity (total-background) of the field of the views were calculated using FIJI software (ImageJ) and plotted against the distance. Control experiments were done without cells in the device, to estimate the diffusion characteristics of the dye through the porous membrane.

#### Pump performance curve

Before starting the experiment, all connectors, tubes and micropipettes were washed with IPA (Sigma W292907), rinsed thoroughly with RO water, dried and then oxygen plasma treated before use. In case of WT cells, after about 3 weeks of confluence, micropipettes were connected to port 3 of the devices and port 0 is closed with a stopper thereby sealing the basal side. This setup was placed inside the incubator such that the micropipette is aligns with the mm scale ruler and the fluid flux in the microcapillary has been recorded at a rate of 60 frames/sec. The height vs time profile was calculated by analyzing the video prepared by stitching the time lapse pictures. The fluid flux was then calculated by multiplying the cross-sectional area of the microcapillary to the slope of the height vs time curve. In case ADPKD cells, micropipettes were connected to port 2 of the devices and port 1 is closed with a stopper thereby sealing the apical side. For fluid shear stress perturbation, a syringe pump was connected to the port 2 and 3 to apply physiological levels of shear stress. For AVP perturbation, the micropipette side was first sealed and then media containing different concentrations of AVP was added and incubated for 30 minutes before starting the time-lapse videography. Similarly, for osmotic perturbation, the micropipette side was first sealed and then hypotonic media was added to the device before starting the time-lapse videography.

#### Immunohistochemistry

Following dissection, tissue samples were fixed in a 4% paraformaldehyde solution, agitated on a rocker for 3-4 hours at 4oC, and then stored at $4^0$C for 24 hours. Following fixation, tissue was washed in 1X PBS, three times for 10-minute washes on a rocker at $4^0$C. Tissue samples were then embedded in paraffin, sectioned, and mounted onto glass coverslips with a gelatin coating solution (50ml distilled water, 0.25g Gelatin, 25mg Chrome Alum). Deparaffination of tissue samples was performed by washing coverslips in a coplin jar with xylene and ethanol (5 minutes in 100% Xylene, 5 minutes in 100% Xylene, 5 minutes in 100% Ethanol, 5 minutes in 95% Ethanol, 5 minutes in 70% Ethanol, 5 minutes in distilled water, 5 minutes in distilled water). Following the last wash, cover slips were placed in a heat induced epitope retrieval (HIER) solution, pH 8.0 (1mM Tris (American Bioanalytical AB02000-01000), 0.5mM EDTA (Sigma E5134) in final volume of 300 mL of distilled water) with 0.02% SDS (American Bioanalytical AB01920-00500). Samples were warmed in HIER solution with SDS to $100^0$C, then transferred to a $100^0$C water bath for 15 minutes. Following this incubation, samples were allowed to cool to room temperature, and washed with distilled water and 1XPBS. Samples were treated with 2-3 drops of Image-iT FX Signal Enhancer (Molecular Probes 136933) for 15 minutes with rotation at room temperature, and then blocked in Incubation Media [1% BSA (Sigma A7638), 0.1% Tween 20 (BioRad 170-6531), 0.02% sodium azide (Sigma S2002) in a final volume of 50 mL 1X PBS (Bio-Rad 161-0780)] with 1% donkey serum (Sigma D9663) for another 15 minutes with rotation at room temperature. Primary antibody was added in incubation media and serum solution [AQP2 (Wade 95 chicken, 1:600) still working on citation] and NKA (Millipore 05-369, 1:200)] and incubated overnight in a humidifier chamber at room temperature. Coverslips were then quickly washed three times in 1X PBS, with a fourth wash lasting for 30 minutes. Secondary antibodies [1:200 Goat anti-chicken secondary antibody (Invitrogen A-21449) and 1:200 Goat anti-mouse secondary antibody (Invitrogen A-11001)] were added in Incubation Media with serum and incubated at room temperature, in the dark for two hours. Cover slips were then again quickly washed (3X) in 1X PBS with a fourth wash lasting 30 minutes. Following washes, cover slips were mounted onto glass slides with Vectashield Mounting Media [Vector Labs H1000], and sealed with nail polish. Samples were imaged on a Olympus IX83 inverted imaging system.

#### Immunofluorescence

Before fixing the both chamber A and chamber B were rinsed gently and thoroughly with PBS (ThermoFisher 14190250). The cells were fixed by adding 3% paraformaldehyde (Electron Microscopy Sciences 15714-S) in PBS in the apical and basal side of the device on ice. After 15 mins the paraformaldehyde was washed off by gently adding PBS three times with a gap of 5 minutes between each wash. The cells were then permeabilized for 10 min susing 0.1% Triton (Sginam T- 9284) in PBS. The cells were then washed with PBS in the same fashion as previously on ice. 5% FBS (ATCC 30-2020) in PBS was used for 30 mins on ice as blocking agent. The cells were then incubated with primary antibodies [AQP2 (Invitrogen PA538004, 1:200) and NaKATPase (Millipore 05-369, 1:200)] with 2.5% FBS in PBS overnight at $40^0$C on a rocker. The primary antibody was washed off using PBS consecutively for three times with 5 minutes interval between each wash. The cells in the device were then incubated with secondary antibodies [1:200 Donkey anti-rabbit secondary antibody (Biolegend 406410) and 1:500 Goat anti-mouse secondary antibody (Invitrogen A-21235)] with 2.5% FBS in PBS for 1 hour at room temperature in the dark. Traces of the secondary antibody was removed from the device by washing the cells with PBS consecutively for three times with an interval of 5 minutes between each wash. F-actin was stained using acti-stain 488 Phalloidin (Cytosckeleton PHDG1-A) at 1:100 with 2.5% FBS in PBS for 1 hour at room temperature. The devices were then filled with Vectashield mounting media (Vector Labs H-1200), sealed with parafilm and stored at $40^0$C.

#### Western Blot

Cystic and non-cystic cells plated on impermeable (regular cell culture dish) and permeable (porous membrane) substrates were extracted for western blot experiments. For lysis, cells were washed twice with PBS and incubates with 100 $\mu$L lysis buffer (25 mM Sodium Phosphate pH7.2, 150 mM NaCl, 10% Glycerol, 1 mM EDTA, 1% Triton, Protease Inhibitor (Roche 11873580001)) for 3 minutes. Plates were scraped and cell solution transferred to 1ml tube. Cell lysates were incubated on ice for 30 mins with regular vortexing. Cell debris was pelleted by centrifugation at 15000 G. samples were prepaired fro SDS-PAGE and equal amounts of protein (75 $\mu$g for each sample) were loaded onto 3-8% Tris-Acetate SDS-polyacrylamide precast gel (invitrogen). Membranes were incubated with primary antibodies overnight, washed with TBS-Tween 20 buffer then incubated with florescent secondary antibodies for detection using a Biorad chemidoc system. Primary antibodies used for immunoblot included mouse anti-PC1 (7e12) from Santa Cruz Biotechnology (sc-130554, 1:500) directed against the LRR; rat monoclonal E8 antibody raised against PC1-CTF (1:1000); Rabbit polyoclonal 3374 antibody raised against C-terminal tail of PC2 (1:1000); mouse anti-$\beta$-actin from Sigma-Aldrich (A5316, 1:10,000).

#### Cystic fluid analysis

Cyst fluid is collected from nephrectomized human ADPKD renal tissue using a 21 gauge needle (BD PrecisionGlide 305165) into a 10 ml latex and silicon oil free syringe (Henk Sass Wolf GMBH 4200.X00V0). Collected fluid is stored at $-80^0$C until time for analysis. Electrolyte analysis is performed using the Easy-Lyte (Medica Corporation, MA), following the manufacturers protocol.

#### Gene expression Analysis

RNA isolation was performed by using the Direct-zolTM RNA MiniPrep Plus (Zymo Research, Irvine, CA) according to manufacturer's instructions. RNA was reverse transcribed by QuantiTect Reverse transcription Kit (QIAGEN, Germantown, MD) and real time PCR was performed by SYBR Green Supermix (Bio-Rad Laboratories, Hercules, CA) using specific primers presented in Table 1. The relative amounts of each mRNA were quantified using the ΔCT method with 18S rRNA as a housekeeping gene. Each experiment was performed two independent times with two or three technical replicates each time. Samples which produced background CT values were discarded. Heatmaps were generated in R by applying the heatmap.2 function and Z-score was calculated using the formula: Z = ($\chi$-$\mu$) / $\sigma$ where $\chi$ is the amount of each mRNA, $\mu$ is the population mean and $\sigma$ is the population standard deviation.

| Gene Name | Forward sequence (5'-3') | Reverse Sequence (5'-3') |
|---|---|---|
| AQP3 | CTCGTGAGCCCTGGATCAAGC | AAAGCTGGTTGTCGGCGAAGT |
| AQP5 | CAGCTGGCACTCTGCATCTT | TGAACCGATTCATGACCACC |
| ATP1A1 | TTAATCCCCCAGGCCTCACT | TCTGGTTATGCCAGAGTGACTG |
| NHE1 | ACCTGGTTCATCAACAAGTTCCG | TTCACAGCCAACAGGTCTACCA |
| SLC12A1 | AGTGCCCAGTAATACCAATCGC | GCCTAAAGCTGATTCTGAGTCTT |
| SLC12A2 | TGGGTCAAGCTGGAATAGGTC | ACCAAATTCTGGCCCTAGACTT |
| CFTR | AGGCAGCCTATGTGAGATAC | TCCGGAGGATGATTCCTTTG |
| TRPM7 | GGAGATGCCCTCAAAGAACA | TGCTCAGGGGGTTCAATAAG |
| TRPV4 | CCCGTGAGAACACCAAGTTT | GTGTCCTCATCCGTCACCTC |
| 18S rRNA | CAGCCACCCGAGATTGAGCA | TAGTAGCGACGGGCGGTGTG |

Table 1: Forward and reverse sequences of primers for the genes used for gene expression analysis

#### Extended figures captions:

##### Extended Data Fig. 1: Fabrication and calibration of Micro-fluidic Kidney Pump (MFKP)

(**a**) Exploded view of the components used in the fabrication and assembly of the Micro-fluidic Kidney Pump (MFKP). The dashed rectangle indicates cross-section of the assembled device with schematic representation of the epithelium grown on the porous membrane. Green arrows indicate fluid flow. (**b**) MFKP was calibrated by measuring the hydrostatic pressure in the apical channel ($P_{\text{apical}}$) while applying fluid shear stress (FSS) on the apical side and changing the hydrostatic pressure in port 2 ($P_2$). During calibration FSS was applied on top of the porous membrane with no cells by changing the flow rate and $P_2$ was increased by increasing the height of the fluid reservoir connected to port 2. Horizontal error bars indicate same-device variation (mean ± standard deviation, n = 3) and vertical error bars indicate device-device variation (mean ± standard deviation, n = 5). Dashed lines indicate values obtained from the FEM simulation under similar velocity and pressure boundary conditions. (**c**) The MC was calibrated using cell culture medium to measure the height of fluid due to capillary action. Error bars indicate standard deviation (n = 5). (**d**) FEM model indicating heat map of the pressure profile in the flow field in MFKP under an FSS of 0.5 $dyn/cm^2$. The arrows indicate direction of fluid flux and the numbers 0,1, 2 and 3 indicate the fluid flow ports. (**e**) Cross-sectional variation of hydrostatic pressure ($P$), fluid velocity magnitude ($v$) and FSS ($\tau_{xy}$) in the XZ plane in MFKP. Apical (A) and basal (B) channels are separated by a porous membrane (bold black line). (**f**) Heatmap indicating the variation of fluid velocity (v) and FSS ($\tau_{xy}$) on top of the porous membrane in MFKP along the XZ plane. (**g**) Heatmap indicating uniform pressure profile along the length of the basal channel (YZ plane) for FSS of both 0.5 $dyn/cm^2$ (**h**) and 1 $dyn/cm^2$ (**i**). The pressure gradient along the apical channel in both FSS is due to the applied flow condition.

##### Extended Data Fig. 2: MDCK-II cells polarize to form strong apical-basal barrier in the MFKP

Confocal fluorescence images of basal and apical channels were taken before and after injection of FITC-conjugated dextran dye of MW 2000 kDa to the apical channel of MFKP in the absence of cells. Average intensity of each slice in the confocal stack were used to quantify the permeability of the porous membrane as well as the barrier strength of the epithelium. (**a**) Without the cells, the intensity of both apical and basal channel is almost zero. However, the average intensity on the basal channel shoots up within a minute of adding the dye to the apical channel. To prevent any leakage the basal channel was closed (by closing ports 0 and 3) before injecting dye into the apical channel. Within 5 minutes the dye diffuses completely into the basal channel. The dashed line indicates the position of the porous membrane that separates the basal and apical channel. (**b**) For MDCK-II epithelium that is 2 days post confluence, it takes about 3 hours for the dye to diffuse into the basal side from the apical side indicating poor barrier strength. (**c**) For MDCK-II epithelium that is 2 weeks post confluence, the dye does not diffuse. The average intensity of the slices in the basal channel remained almost same after 3 hours post injection of dye in the apical channel. (**d**) Comparison of fluorescence images of the basal and apical channel in MFKP with and without the dye (green) indicate diffusion of the dye through the porous membrane. The schematic on the left demonstrates experimental condition and green arrows indicate the direction of diffusion of the dye. (**e**) For an MDCK-II epithelium that is 2 days post confluence, representative images of the basal and apical channel in MFKP at T = 0 hour and at T = 3 hours indicate slow diffusion of the dye through the epithelium. (**f**) For an MDCK-II epithelium that is 2 weeks post confluence, representative images of the basal and apical channel in MFKP at T = 0 hour and at T = 3 hours indicate no dye diffusion through the epithelium.

#### Extended Data Fig. 3: MDCK-II domes exhibit similar trans-epithelial hydrostatic pressure gradient

(**a**) Brightfield image showing a micro-needle inserted into a fluid-filled MDCK-II dome formed on glass substrate. Scale bar = 25 $\mu$m. (**b**) Schematic of micro-needle inserted into a fluid filled epithelial dome. Asterisk indicates lumen side in the dome. (**c**) Dashed rectangle shows phase-contrast image of oil-media interface in the micro-needle. The curvature of the interface was used to measured the hydrostatic pressure in the domes. $P_2$ and $P_1$ indicate the hydrostatic pressure at the oil-media interface in the micro-needle. Scale bar = 20 $\mu$m. (**d**) Distribution of fluid-filled dome sizes in an MDCK-II epithelium on glass. (**e**) Distribution of hydrostatic pressure measured in fluid-filled domes in an MDCK-II epithelium on glass. Error bars indicate standard deviation. Fluid-filled domes on impermeable substrates were used to investigate the effect of stall pressure on the colocalization of F-actin and NKA in MDCK-II cells. (**f**) Schematic representation of an unstable MDCK-II dome, wherein the epithelium has just started to accumulate fluid on the basal side. This situation is $\Delta P \approx 0$. (**g**) Schematic representation of a stable MDCK-II dome, where the hydrostatic pressure gradient approximately equal to the stall pressure ($\Delta P^*$). (**h**) Confocal reconstruction of an unstable dome showed colocalization of F-actin (green) and NKA (red) in the cells. Scale bar = 10$\mu$m. (**i**) Zoomed view of a cell indicated by dashed rectangle in **h**, showing enrichment of F-actin in the cortex. (**j**) Zoomed view of the cell indicated by dashed rectangle in **h**, showing enrichment of NKA on the basolateral domain. Scale bar = 10$\mu$m. (**k**) Intensity profiles of F-actin (green) and NKA (red) along the band in **i** and **j**. The pink rectangle indicates location of cell with the band and A and B indicate apical and basal domain of the cell under consideration. The asterisk indicates basal side of the cells. (**l**) Confocal reconstruction of a stable dome showing colocalization of F-actin (green) and NKA (red) in the cells. Scale bar = 10$\mu$m. (**m**) Cross-sectional view of the dome indicated by dashed line in **l**, showing enrichment of F-actin in the cells. **n**, Cross-sectional view of the dome indicated by dashed line in **l**, showing enrichment of NKA in the cells. Scale bar=10$\mu$m. (**o**) Intensity profile of F-actin (green) and NKA (red) along the band in **m** and **n**. (**p**) Comparison of the total intensity of F-actin along the cell height in five cells chosen arbitrarily in the two conditions: unstable domes ($\Delta P \approx 0$) and stable domes ($\Delta P \approx \Delta P^*$). (**q**) Comparison of the total intensity of NKA along the cell height in five cells chosen arbitrarily in the two conditions: unstable domes ($\Delta P \approx 0$) and stable domes ($\Delta P \approx \Delta P^*$). Purple triangles indicate ($\Delta P \approx 0$) and black circles indicates ($\Delta P \approx \Delta P^*$).

#### Extended Data Fig. 4: Phenotypic resemblance of both Wild Type (WT) and ADPKD epithelium grown in MFKP to that of human kidney sections

(**a**) Immunohistochemical analysis of tissue section of normal human kidney and ADPKD kidneys reveal that in case of both normal tubules and cysts, Na/K ATPase (NKA, red) is located primarily on the basolateral side. (**b**) Intensity profile of NKA along the lines shown in **a**. The pink line plot is the NKA intensity profile along the pink line in a non-cystic cell. The blue line plot is the NKA intensity profile along the blue line on a cystic cell. A and B mark the apical and basal surface of cells under consideration. (**c**) Aquaporin -2 (AQP2, blue) is however located on the apical or sometimes sub-apical domain of cells in both normal human kidney tissue sections and cystic section in APPKD kidneys. (**d**) Intensity profile of AQP2 along the lines shown in

**c**. The green line plot is the AQP2 intensity profile along the green line on a non-cystic cell. The orange line plot is the AQP2 intensity profile along the orange line on a cystic cell. The asterisk indicates the lumen side in cystic tissue section. Both NKA and AQP2 stains indicate that the apico-basal polarity remains the same both in normal tubules and cysts, where the apical side is facing the lumen and the basal side is facing the interstitial. Normal cells demonstrate regular cuboidal columnar morphology. However, the cystic cells have irregular and stretchy morphology coupled with decreased cell height. (**e**) Plot showing the concentration of $Na^+$, $K^+$ and $Cl^-$ and osmolality of the cystic fluid from ADPKD patients. (**f**), (**h**) and (**j**) Confocal immunofluorescence images of WT cortex (WTc), WT medulla (WTm) and ADPKD cells extracted from normal human kidneys and ADPKD kidneys seeded in MFKP, showing co-localization of AQP2 (blue), NKA (red) and F-actin (green). The dashed white line in XZ images represents the porous membrane. (**g**) Quantification of apico-basal intensity for AQP2, NKA and F-actin in WTc epithelium grown on MFKP shows enrichment of NKA primarily in the basolateral domain of the cells indicated by white arrow on the XZ image. AQP2 signal was low indicating absence of the protein in WTc cells. F-actin stress fibers were present in the basal side and strong enrichment in the cell-cell junctions (indicated by arrow) was observed. F-actin XZ images demonstrate that WTc epithelium exhibit the regular columnar cuboidal epithelium. (**i**) In case of WTm cells, AQP2 was highly enriched in the apical domain and is also present all over the cytoplasm. NKA intensity was low and is primarily located in the basal side of the cells. The F-actin is fibrous throughout the cytoplasm and strong enrichment in the cell-cell junctions was observed (marked by arrow in the F-actin XZ image. (**k**) ADPKD cells showed AQP2 expression on the apical side in a sporadic fashion in the epithelium. NKA is enriched in the basolateral domain and is not uniformly distributed all over the epithelium. F-actin is highly fibrous all over the cytoplasm and unlike normal cells, localization in the cell-cell junction was relatively low. Scale bar=50$\mu$m.

#### Extended Data Fig. 5: Normal human kidney tubular epithelial and ADPKD cystic cells polarize to form strong apico-basal barrier in the MFKP

Confocal fluorescence images of the basal and the apical channel were taken before and after injection of FITC-conjugated dextran dye of MW 2000 kDa into the apical channel of MFKP in the absence of cells. Average intensity of the stack of images were used to quantify the permeability of the porous membrane as well as the barrier strength of the epithelium. (**a**) For WTc epithelium after 2 weeks post confluence, the dye did not diffuse from the apical to basal side. The average dye intensity of the basal channel remained the same 4 hours post injection of dye in the apical channel. The vertical dashed line (black) is the position of the porous membrane that separates the basal and apical side. (**b**) For WTm epithelium after 2 weeks post confluence, the dye did not diffuse from the apical to basal side. The average dye intensity of the basal channel remained the same 4 hours post injection of dye in the apical channel.(**c**) In case of ADPKD epithelium after 2 weeks post confluence, the dye did not diffuse from the apical to basal side. The average dye intensity of the basal channel remained the same 4 hours post injection of dye in the apical channel. Representative fluorescence images of basal and apical channel at T = 0 hour and at T = 4 hours for WTc (**d**), WTm (**e**) and ADPKD cells (**f**). The horizontal dashed line (white) is the position of the porous membrane that separates the basal and apical side. The variability of the average intensity values in the same device and from device to device is plotted for WTc (**g**), WTm (**h**) and ADPKD cells (**i**). Horizontal error bars indicate same-device variability and vertical error bars indicate device-to-device variability (mean $\pm$ standard deviation, n = 3).

##### Extended Data Fig. 6: Fluid pumping performance curve for WTc, WTm and ADPKD cells are dependent on mechanical, chemical and osmotic perturbations

(**a**) Pump performance curve (PPC) shows that the trans-epithelial fluid flux (J) for wild type cortical cells (WTc) decreases with increase in hydrostatic pressure ($\Delta P$) across the epithelium. With 20 % hypotonic osmolarity change in the apical channel, $J_0$ went up to 8.5 $\mu$L/min/cm$^2$ and $\Delta P^*$ to 230 Pa. Basolateral treatment with 10 nM Argnine Vasopressin (AVP) did not change the $J_0$, but lowered $\Delta P^*$. (**b**) In case of wild type medulla cells (WTm), the PPC was much steeper with $J_0$ = 7 $\mu$L/min/cm$^2$ and $\Delta P^*$ = 60 Pa. With 20 % hypotonic osmolarity change in the apical channel, both $J_0$ and $\Delta P^*$ went up. Unlike WTc cells, basolateral perturbation by 10 nM AVP caused both $J_0$ and $\Delta P^*$ to increase. (**c**) In case of ADPKD cells, cells pump fluid from the basal to apical channel therefore developing a stall pressure $\Delta P^*$ = -300 Pa. With basal hypo-osmotic perturbation of 20%, the $J_0$ value went up to 9 $\mu$L/min/cm$^2$ as compared to 3$\mu$L/min/cm$^2$ for control cells. ADPKD cells when treated with 20 nM AVP basolaterally, caused increase in $J_0$ to 14 $\mu$L/min/cm$^2$. However, under both osmotic gradient and AVP perturbations, the stall pressure $\Delta P^*$ did not change. (**d**) Schematic of the theoretical model of fluidic pumping by epithelial cells. Flow field along the $x$-direction and apical and basal surface water and solute fluxes are considered by the model. (**e**) Theoretical pump performance curve calculated for the model epithelium using active flux in Eq. (19), where $J$ is the trans-epithelial fluid flux and $\Delta P$ is the hydrostatic pressure gradient. (**f**) Converging nature of PPC to a constant $\Delta P^*$ was achieved by changing the $J^{\mathrm{a}}_{\mathrm{active}}$ to Eq. (20). Lighter lines indicate decrease in apical osmolarity. The fluid flux values were normalized against the maximum $J_0$.

##### Extended Data Fig. 7: Differential ion channel expression of normal and diseased cells grown in MFKP as compared to impermeable substrates.

(**a**) Heatmaps showing relative expressions of mRNAs extracted from WTc, WTm and ADPKD cells grown on permeable substrate (MFKP) and on impermeable substrate (tissue culture treated polystyrene dishes). P and I indicate permeable and impermeable substrate, respectively. The rows are normalized such that the relative concentration across the cells lines has been shown using color code. The list of genes includes Aquaporins (AQP1, AQP3 and AQP5) and ion-pump (NKA) and ion-channels (NHE1, NKCC1, NKCC2, CFTR, TRPM7 and TRPV4). Two biological repeats were done for each condition. (**b**) Western blot with an antibody directed against PC1 C-terminus (top panel). Total lysate of cells derived from human WT cortex (WTc), WT medulla (WTm) and ADPKD kidneys grown on impermeable and permeable (MFKP) substrates were loaded on the gel. PC1 full length (PC1 FL) and PC1 CTF and PC1 NTF bands are indicated. Below, anti-PC2 western blot shows PC2 band at ~110 kDa. Densitometry of relative protein abundance of each band is represented graphically for PC-1 (**c**), PC 1 FL (**d**), PC1 NTF (**e**), PC2 (**f**).

##### Extended Data Fig. 8: Role of stall pressure on NKA in WTc cells.

(**a-b**) Comparison of $J_0$ and $\Delta P^*$ for WTc epithelium in control and 274 $\mu M$ Ouabain treatment for 12 hours. (**c**) XZ confocal section of WTc epithelium in MFKP showing localization of F-actin (green), NKA (red) in the cells at zero trans-epithelial hydrostatic pressure gradient (or $\Delta P$ = 0). The white dashed line represents the porous membrane. (**d**) Normalized intensity profiles of F-actin (green) and NKA (red) along the white band in **c**. The pink rectangle indicates location of the cell under consideration. A and B indicate apical and basal domains of the cell under consideration. (**e**) XZ confocal section of WTc epithelium in MFKP showing localization of F-actin (green), NKA (red) in the cells at stall pressure (or $\Delta P$ = $\Delta P^*$

$\approx$ 200 Pa). (**f**) Normalized intensity profiles of F-actin (green) and NKA (red) along the band in **e**. The pink rectangle indicates location of the cell under consideration in **b**. (**g**) Maximum intensity projection of the 40 $\mu$m thick confocal stacks (41 sections at 1 $\mu$m intervals) of F-actin at $\Delta P$ = 0. (**h**) Maximum intensity projection of the 40 $\mu$m thick confocal stacks (41 sections at 1 $\mu$m intervals) of NKA ($\alpha$ sub-unit) at $\Delta P$ = 0. (**i**) Maximum intensity projection of the 40 $\mu$m thick confocal stacks (41 sections at 1 $\mu$m intervals) of F-actin at $\Delta P$ = $\Delta P$*. (**j**) Maximum intensity projection of the 40 $\mu$m thick confocal stacks (41 sections at 1 $\mu$m intervals) of NKA ($\alpha$ sub-unit) at $\Delta P$ = $\Delta P$*. (**k**) Zoomed IF image (represented by the dashed square in **g**) of a single slice showing cortical F-actin in WTc cells at $\Delta P$ = 0. (**l**) Zoomed IF image (represented by the dashed square in **h**) of a single stack showing cortical NKA in WTc cells at $\Delta P$ = 0. (**m**) Zoomed IF image (represented by the dashed square in **i**) of a single slice showing cortical F-actin in WTc cells at $\Delta P$ = $\Delta P$*. (**n**) Zoomed IF image (represented by the dashed square in **j**) of a single stack showing cortica NKA in WTc cells at $\Delta P$ = $\Delta P$*. (**o**) Comparison of the total intensity of F-actin in six cells chosen arbitrarily in **g** and **i** is plotted along the height of the cells. Letters B and A in the plot indicate the location of basal and apical surface of cells. Purple triangles indicate $\Delta P$ = 0 and black circles indicates $\Delta P$ = $\Delta P$*. (**p**) Comparison of the total intensity NKA in five cells chosen arbitrarily in **h** and **j** is plotted along the height of the cells. Purple triangles indicate $\Delta P$ = 0 and black circles indicates $\Delta P$ = $\Delta P$*. (**q**) Comparison of the cortical intensities of F-actin at $\Delta P$ = 0 and $\Delta P$ = $\Delta P$*. The blue line is the cortical intensity profile of F-actin in case of $\Delta P$ = 0 along the band (blue) in **k**. The pink line is the cortical intensity profile of F-actin in case of $\Delta P$ = $\Delta P^*$ along the band (pink) in **m**. (**r**) Comparison of the cortical intensities of NKA at $\Delta P$ = 0 and $\Delta P$ = $\Delta P$*. The blue line is the cortical intensity profile of NKA in case of $\Delta P$ = 0 along the band (blue) in **l**. The pink line is the cortical intensity profile of NKA in case of $\Delta P$ = $\Delta P^*$ along the band (pink) in **n**. (**s**) Schematic representation of the spatial localization of F-actin (green lines) and NKA (red dots) in WTc cells in MFKP at $\Delta P$ = 0. (**t**) Schematic representation of the spatial localization of F-actin (green lines) and NKA (red dots) in WTc cells in MFKP at $\Delta P$ = $\Delta P$*. Scale bar=20$\mu$m.

#### Extended Data Fig. 9: Role of stall pressure on NKA in WTm cells.

(**a-b**) Comparison of $J_0$ and $\Delta P^*$ for WTm epithelium in control and 274 $\mu$M Ouabain treatment for 12 hours. (**c**) XZ confocal section of WTm epithelium in MFKP showing localization of F-actin (green), NKA (red) in the cells at zero trans-epithelial hydrostatic pressure gradient (or $\Delta P$ = 0). The white dashed line represents the porous membrane. (**d**) Normalized intensity profiles of F-actin (green) and NKA (red) along the white band in **c**. The pink rectangle indicates location of the cell under consideration. (**e**) XZ confocal section of WTm epithelium in MFKP showing localization of F-actin (green), NKA (red) in the cells at stall pressure (or $\Delta P$ = $\Delta P^*$ $\approx$ 200 Pa). (**f**) Normalized intensity profiles of Factin (green) and NKA (red) along the band in **b**. The pink rectangle indicates location of the cell under consideration in **e**. (**g**) Maximum intensity projection of the 40 $\mu$m thick confocal stacks (41 sections at 1 $\mu$m intervals) of F-actin at $\Delta P$ = 0. (**h**) Maximum intensity projection of the 40 $\mu$m thick confocal stacks (41 sections at 1 $\mu$m intervals) of NKA ($\alpha$ sub-unit) at $\Delta P$ = 0. (**i**) Maximum intensity projection of the 40 $\mu$m thick confocal stacks (41 sections at 1 $\mu$m intervals) of F-actin at $\Delta P$ = $\Delta P$*. (**j**) Maximum intensity projection of the 40 $\mu$m thick confocal stacks (41 sections at 1 $\mu$m intervals) of Na/K ($\alpha$ sub-unit) at $\Delta P$ = $\Delta P$*. (**k**) Zoomed IF image (represented by the dashed square in **g**) of a single stack showing cortical F-actin in WTm cells at $\Delta P$ = 0. (**l**) Zoomed IF image (represented by the dashed square in **h**) of a single stack showing cortica NKA in WTm cells at $\Delta P$ = 0. (**m**) Zoomed IF image (represented by the dashed square in **i**) of a single stack showing cortical F-actin in WTm cells at $\Delta P$ = $\Delta P$*. (**n**) Zoomed IF image (represented by the dashed square in **j**) of a single stack showing cortical NKA in WTm cells at $\Delta P$ = $\Delta P^{*}$. (**o**) Comparison of the total intensity of F-actin in six cells chosen arbitrarily in **g** and **i** is plotted along the height of the cells. Letters B and A in the plot indicate the location of basal and apical surface. Purple triangles indicate $\Delta P$ = 0 and black circles indicates $\Delta P$ = $\Delta P^{*}$. (**p**) Comparison of the total intensity NKA in five cells chosen arbitrarily in **h** and **j** is plotted along the height of the cells. Purple triangles indicate $\Delta P$ = 0 and black circles indicates $\Delta P$ = $\Delta P^{*}$. (**q**) Comparison of the cortical intensities of F-actin at $\Delta P$ = 0 and $\Delta P$ = $\Delta P^{*}$. The blue line is the cortical intensity profile of F-actin in case of $\Delta P$ = 0 along the band (blue) in **k**. The pink line is the cortical intensity profile of F-actin in case of $\Delta P$ = $\Delta P^{*}$ along the band (pink) in **m**. (**r**) Comparison of the cortical intensities of NKA at $\Delta P$ = 0 and $\Delta P$ = $\Delta P^{*}$. The blue line is the cortical intensity profile of NKA in case of $\Delta P$ = 0 along the band (blue) in **l**. The pink line is the cortical intensity profile of NKA in case of $\Delta P$ = $\Delta P^{*}$ along the band (pink) in **n**. (**s**), Schematic representation of spatial localization of F-actin (green lines) and NKA (red dots) in WTm cells in MFKP at $\Delta P$ = 0. (**t**) Schematic representation of the spatial arrangement of F-actin (green lines) and NKA (red dots) in WTm cells in MFKP at $\Delta P$ = $\Delta P^{*}$. Scale bar=20$\mu$m.

##### Extended Data Fig. 10: Role of stall pressure on NKA in ADPKD cells.

(**a-b**) Comparison of $J_0$ and $\Delta P^{*}$ for ADPKD epithelium in control and 274 $\mu$M Ouabain treatment for 12 hours. (**c**) XZ confocal section of ADPKD epithelium in MFKP showing colocalization of F-actin (green), NKA (red) in the cells at zero trans-epithelial hydrostatic pressure gradient (or $\Delta P$ = 0). The white dashed line represents the porous membrane. (**d**) Normalized intensity profiles of F-actin (green) and NKA (red) along the white band in **c**. The pink rectangle indicates location of the cell under consideration. (**e**) XZ confocal section of ADPKD epithelium in MFKP showing localization of F-actin (green), NKA (red) in the cells at stall pressure (or $\Delta P$ = $\Delta P^{*}$ $\approx$ 200 Pa). (**f**) Normalized intensity profiles of F-actin (green) and NKA (red) along the band in **e**. The pink rectangle indicates location of the cell under consideration. (**g**) Maximum intensity projection of the 40 $\mu$m thick confocal stacks (41 sections at 1 $\mu$m intervals) of F-actin at $\Delta P$ = 0. (**h**) Maximum intensity projection of the 40 $\mu$m thick confocal stacks (41 sections at 1$\mu$m intervals) of NKA ($\alpha$ sub-unit) at $\Delta P$ = 0. (**i**) Maximum intensity projection of the 40 $\mu m$ thick confocal stacks (41 sections at 1 $\mu m$ intervals) of F-actin at $\Delta P$ = $\Delta P^{*}$. (**j**) Maximum intensity projection of the 40 $\mu$m thick confocal stacks (41 sections at 1$\mu$m intervals) of Na/K ($\alpha$ sub-unit) at $\Delta P$ = $\Delta P^{*}$. (**k**) Zoomed IF image (represented by the dashed square in **g**) of a single stack showing cortical F-actin in ADPKD cells at $\Delta P$ = 0. (**l**) Zoomed IF image (represented by the dashed square in **h**) of a single stack showing cortical NKA in WTc cells at $\Delta P$ = 0. (**m**) Zoomed IF image (represented by the dashed square in **i**) of a single stack showing cortical F-actin in ADPKD cells at $\Delta P$ = $\Delta P^{*}$. (**n**) Zoomed IF image (represented by the dashed square in **j**) of a single stack showing cortica NKA in ADPKD cells at $\Delta P$ = $\Delta P^{*}$. (**o**) Comparison of the total intensity of F-actin in six cells chosen arbitrarily in **g** and **i** is plotted along the height of the cells. B and A i indicate the location of basal and apical surfaces of cells. Purple triangles indicate $\Delta P$ = 0 and black circles indicates $\Delta P$ = $\Delta P^{*}$. (**p**) Comparison of the total intensity NKA in five cells chosen arbitrarily in **h** and **j** is plotted along the height of the cells. Purple triangles indicate $\Delta P$ = 0 and black circles indicates $\Delta P$ = $\Delta P^{*}$. (**q**) Comparison of the cortical intensities of F-actin at $\Delta P$ = 0 and $\Delta P$ = $\Delta P^{*}$. The blue line is the cortical intensity profile of F-actin in case of $\Delta P$ = 0 along the band (blue) in **k**. The pink line is the cortical intensity profile of F-actin in case of $\Delta P$ = $\Delta P^{*}$ along the band (pink) in **m**. (**r**) Comparison of the cortical intensities of NKA at $\Delta P$ = 0 and $\Delta P$ = $\Delta P^{*}$. The blue line is the cortical intensity profile of NKA in case of $\Delta P$ = 0 along the band (blue) in **l**. The pink line is the cortical intensity profile of NKA in case of $\Delta P$ = $\Delta P^*$ along the band (pink) in **n**. (**s**) Schematic representation of the localization of F-actin (green lines) and NKA (red dots) in ADPKD cells in MFKP at $\Delta P$ = 0. **t**, Schematic representation of the localization of F-actin (green lines) and NKA (red dots) in ADPKD cells in MFKP at $\Delta P$ = $\Delta P^*$. Scale bar=20$\mu$m.

#### Video Captions:

- Video 1: Time-lapsed videography of the fluid pumping action of MDCK-II cells in MFKP and corresponding pump performance curve

  Time lapsed videography of the fluid pumping action of MDCK-II epithelium grown in MFKP. Plot representing the variation of fluid height in the MC with time. The pump performance curve of WTc epithelium indicating the variation of trans-epithelial fluid flux $(J)$ with the hydrostatic pressure gradient ($\Delta P$) developed across the epithelium. Time step is 1 min.

- Video 2: Time-lapsed videography of the fluid pumping action of WTc, WTm and ADPKD cystic cells in MFKP and corresponding pump performance curve

  Time lapsed videography of the fluid pumping action of WTc, WTm and ADPKD epithelium grown in MFKP. Variation of fluid height in the MC with time for WTc, WTm and ADPKD cells. The pump performance curve of WTc, WTm and ADPKD cells indicating the variation of trans-epithelial fluid flux $(J)$ with the hydrostatic pressure gradient ($\Delta P$) developed across the epithelium. Time step is 1 min.

- Video 3: Fluctuating epithelial dome

  Differential interference contrast (DIC) time-lapsed video of an unstable fluctuating MDCK-II domes on glass substrate. Original time step is 30 mins. Video was saved at 5 frames per second. scale bar = 50 $\mu$m

- Video 4: Mature epithelial dome

  Differential interference contrast (DIC) time-lapsed video of stable MDCK-II domes on glass substrate. Original time step is 30 mins. Video was saved at 5 frames per second. scale bar = 50 $\mu$m.

Extended Figure 1

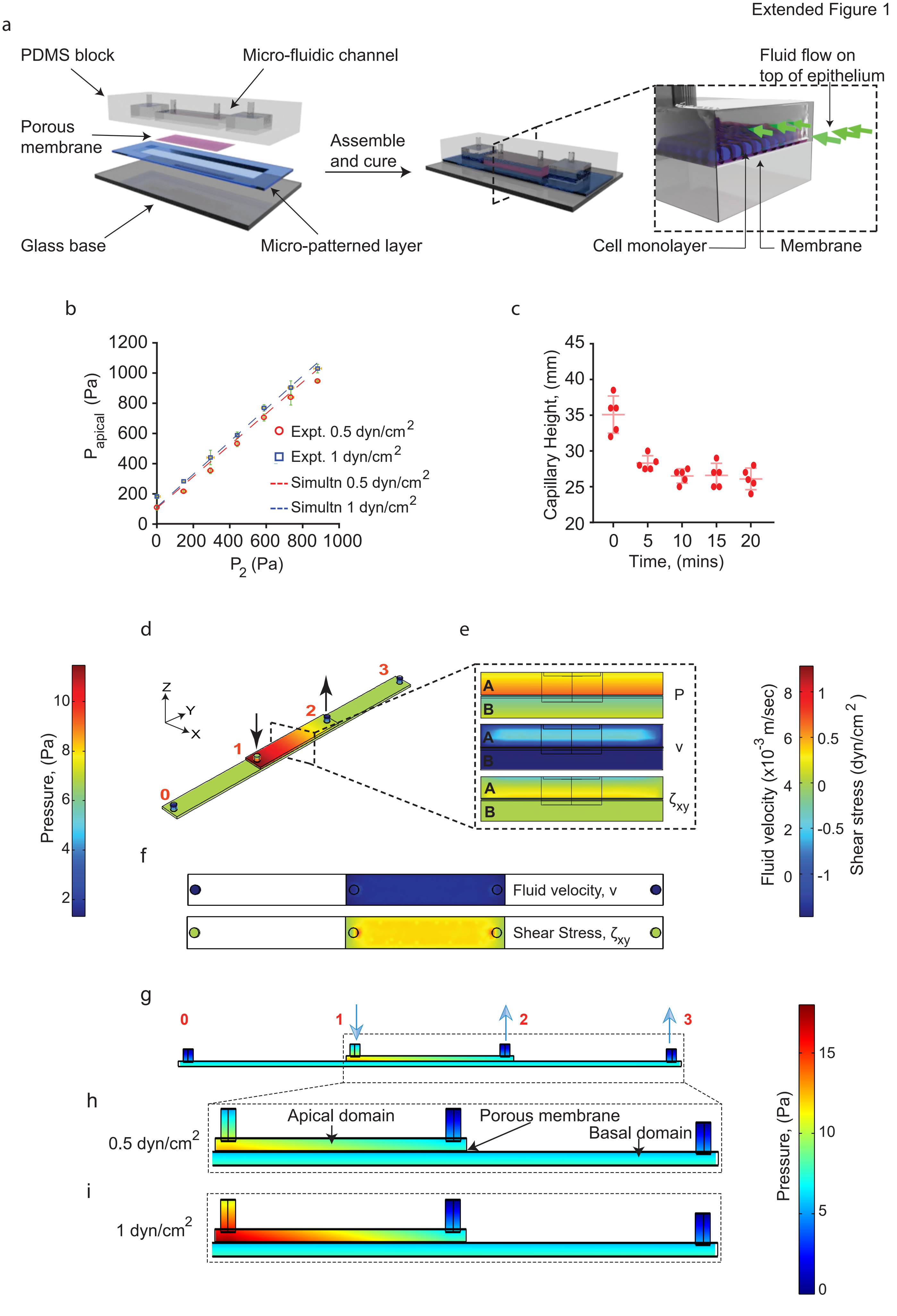

Extended Figure 2

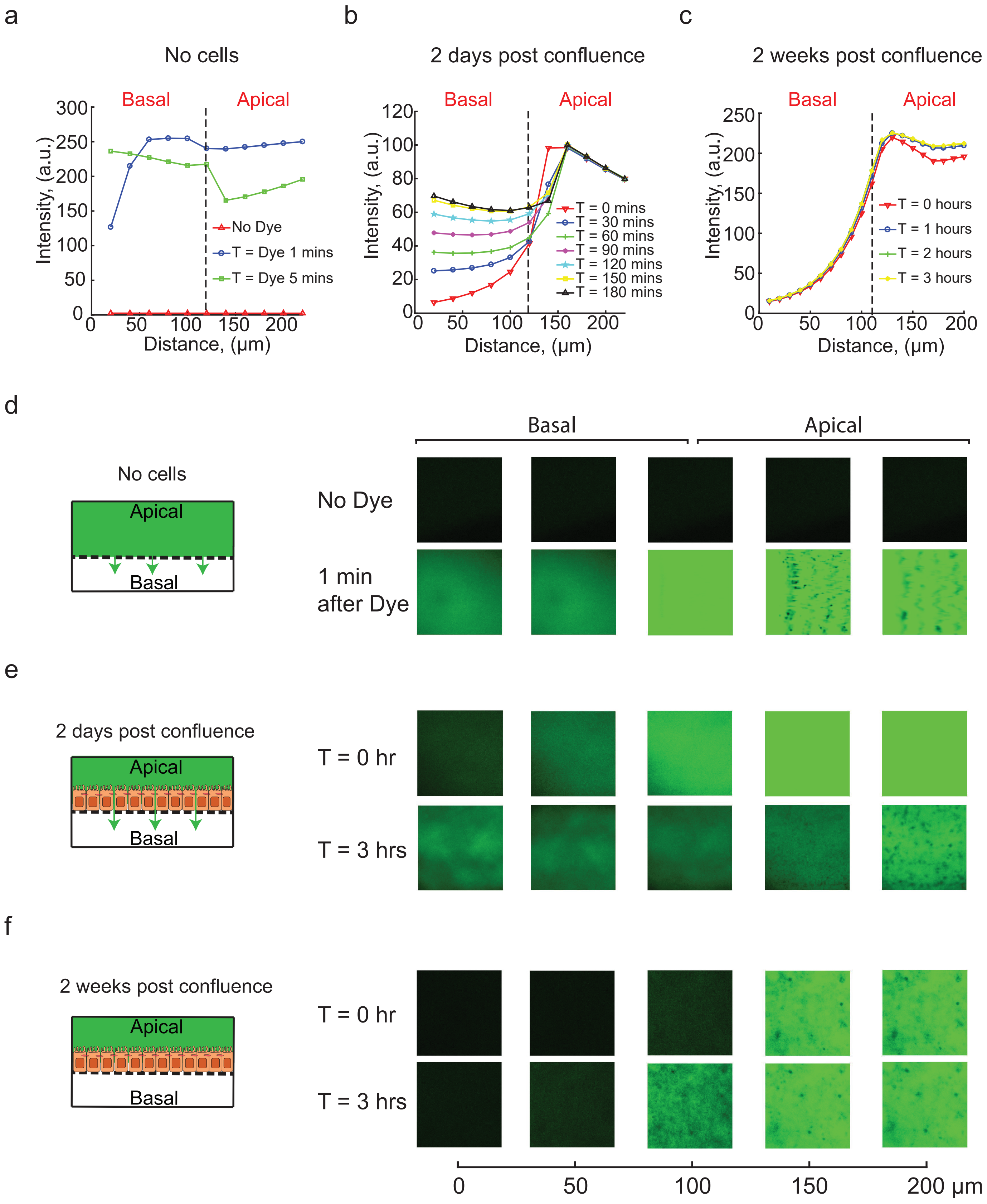

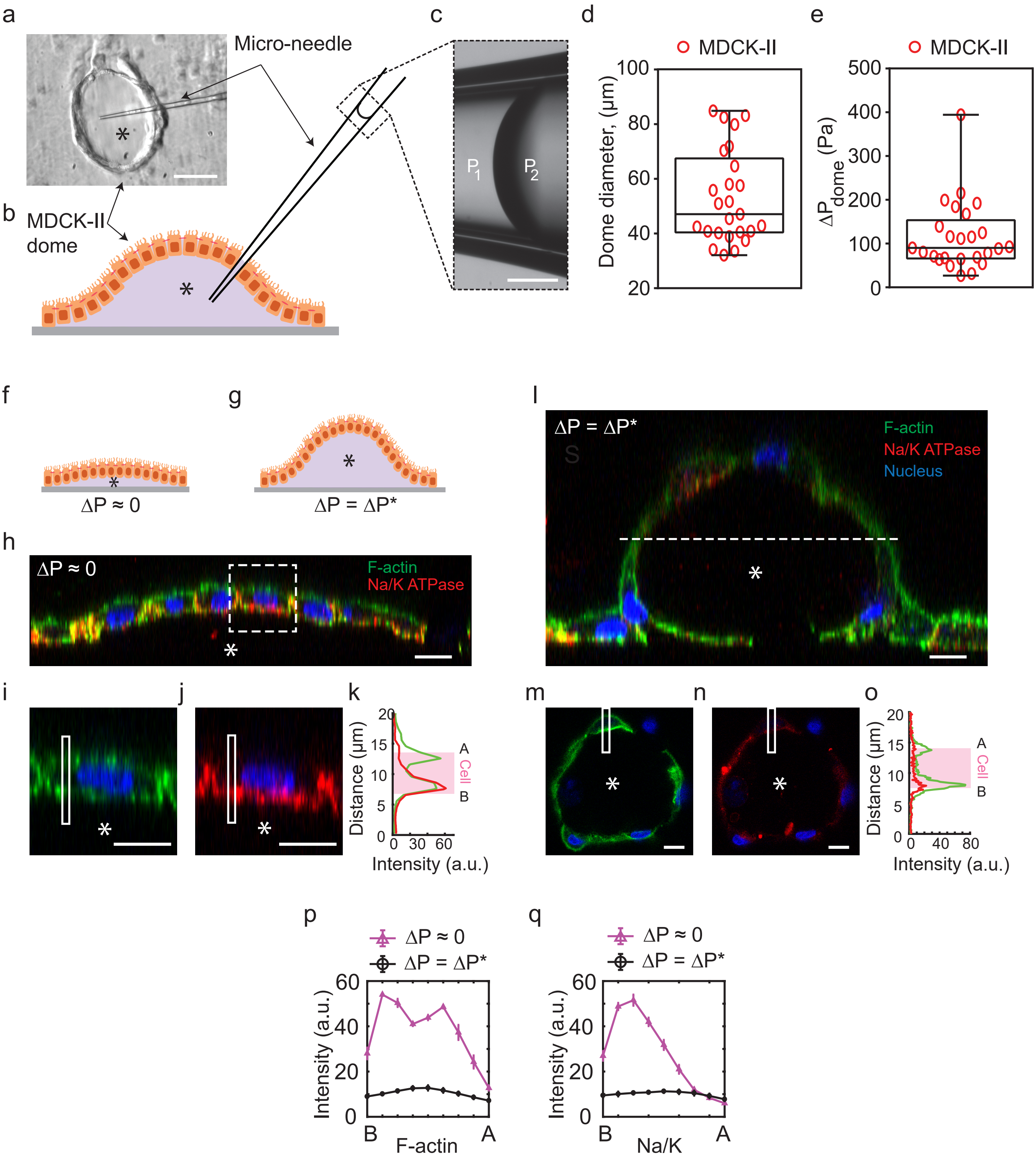
a
Micro-needle
b
MDCK-II
dome
c
P1
P2
d
MDCK-II
Dome diameter, (µm)
e
MDCK-II
ΔPdome (Pa)
f
ΔP ≈ 0
g
ΔP = ΔP*
h
ΔP ≈ 0
F-actin
Na/K ATPase
l
ΔP = ΔP*
F-actin
Na/K ATPase
Nucleus
i
j
k
Distance (µm)
Intensity (a.u.)
Cell
A
B
m
n
o
Distance (µm)
Intensity (a.u.)
Cell
A
B
p
ΔP ≈ 0
ΔP = ΔP*
Intensity (a.u.)
F-actin
B
A
q
ΔP ≈ 0
ΔP = ΔP*
Intensity (a.u.)
Na/K
B
A

Extended Figure 4

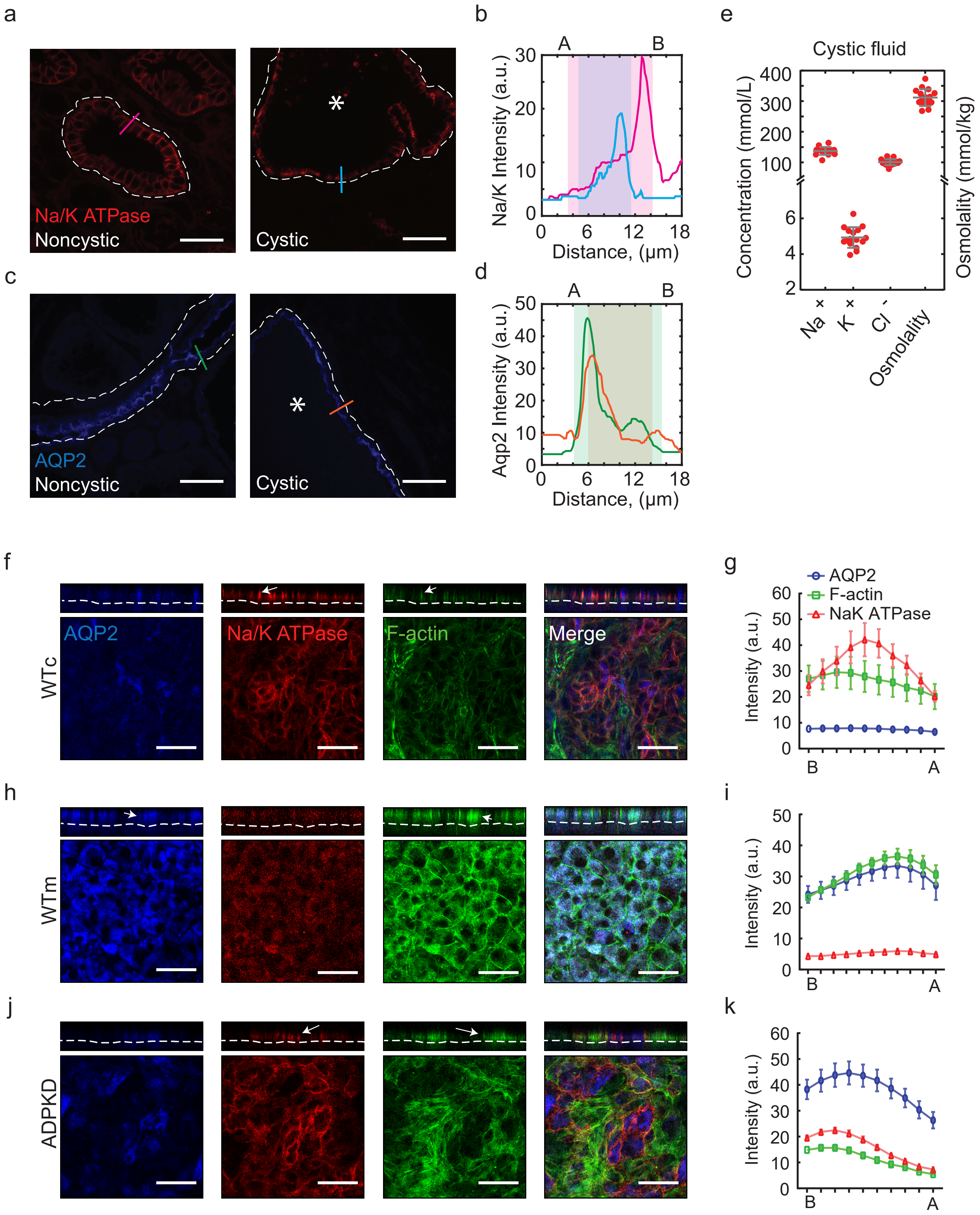

Extended Figure 5

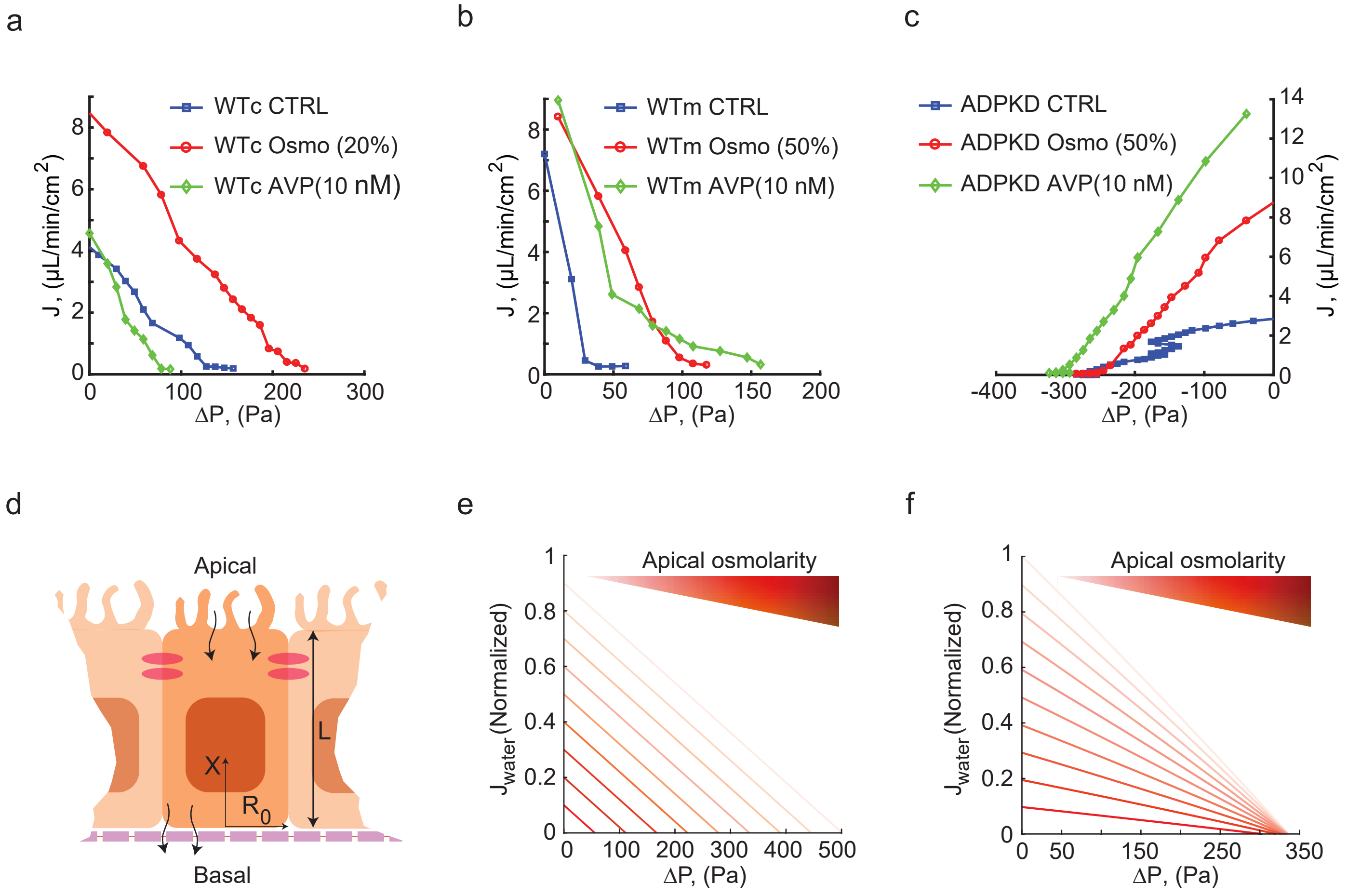

Extended Figure 6

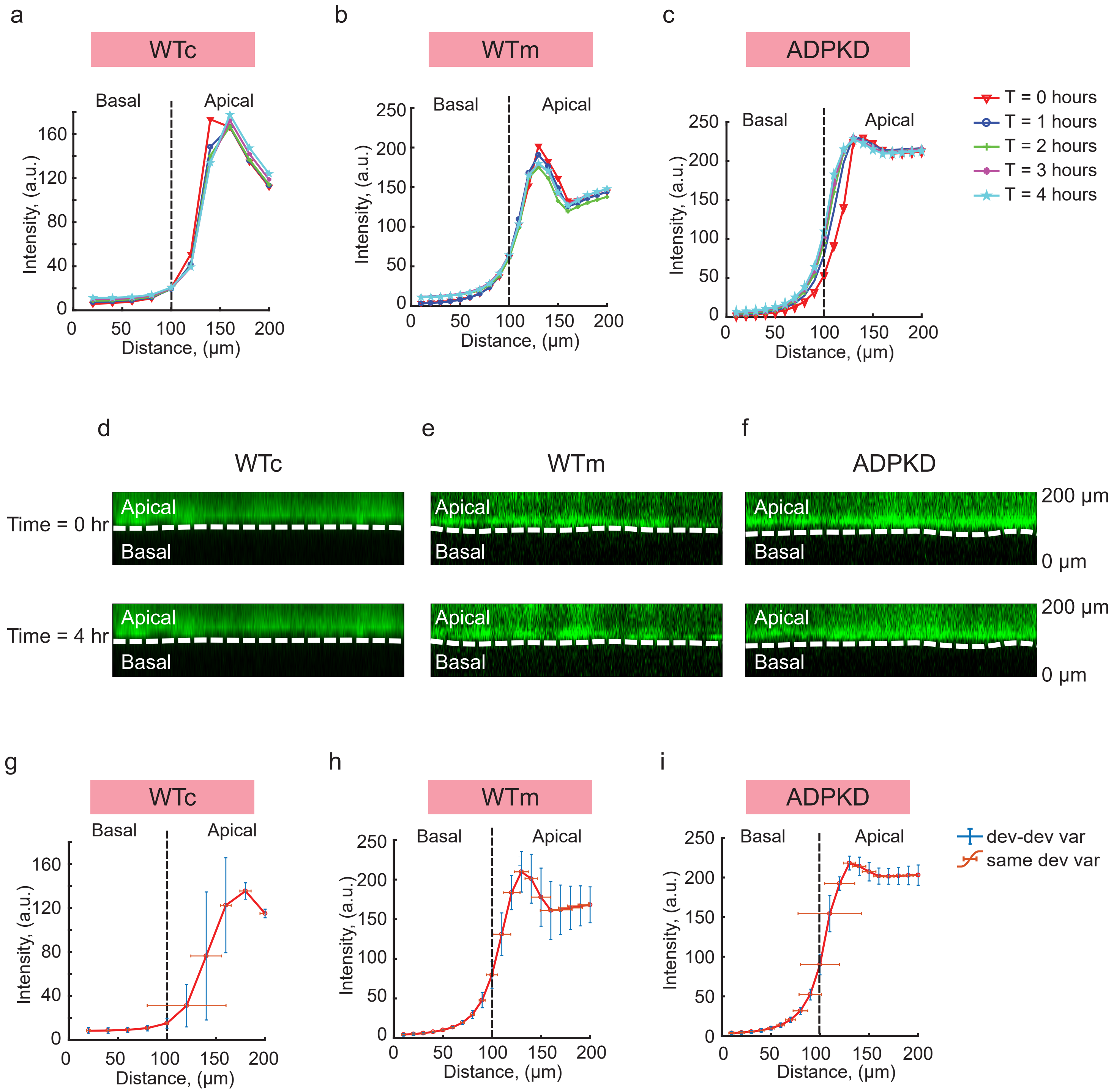

Extended Figure 7

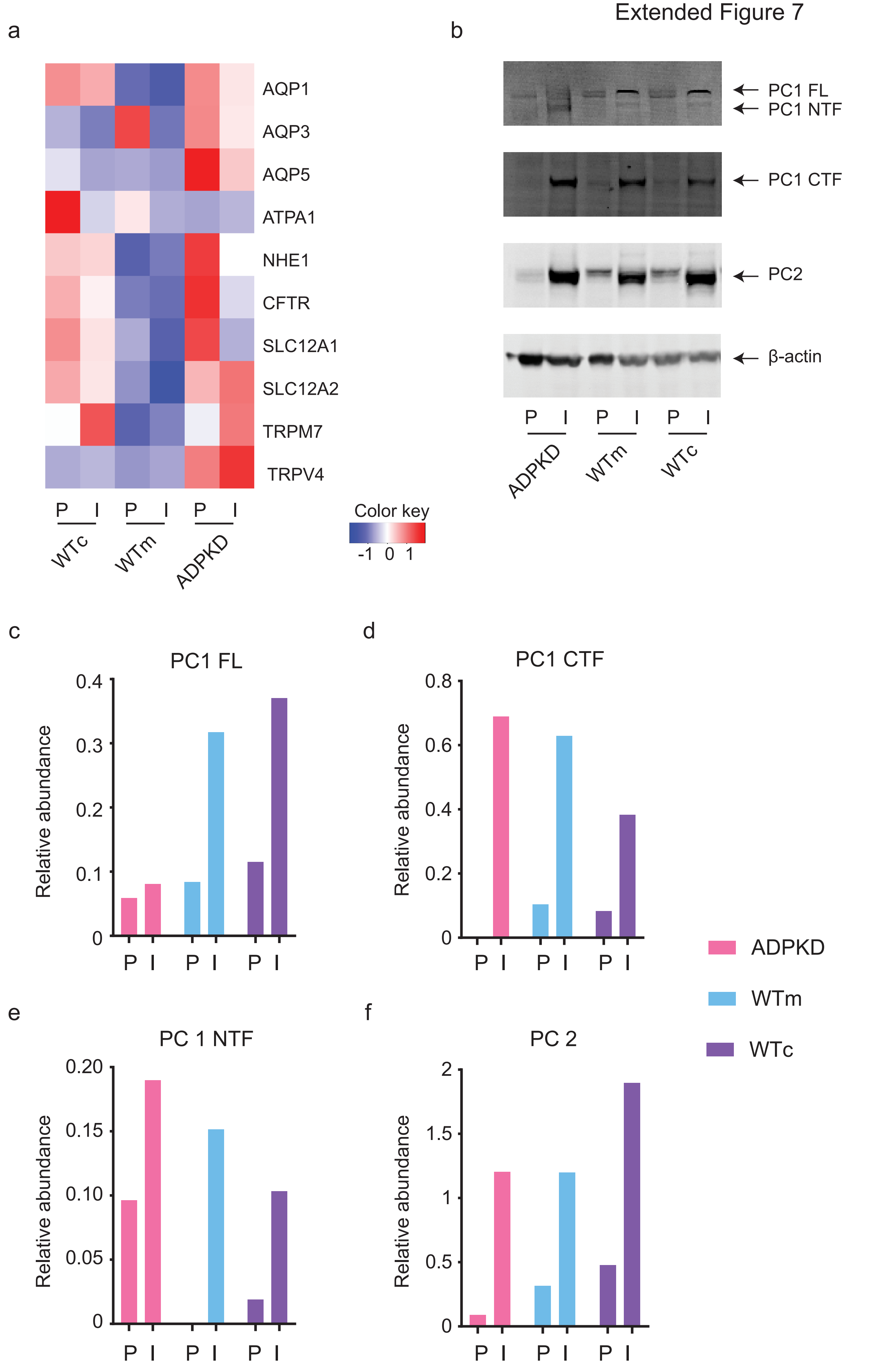

Extended Figure 8

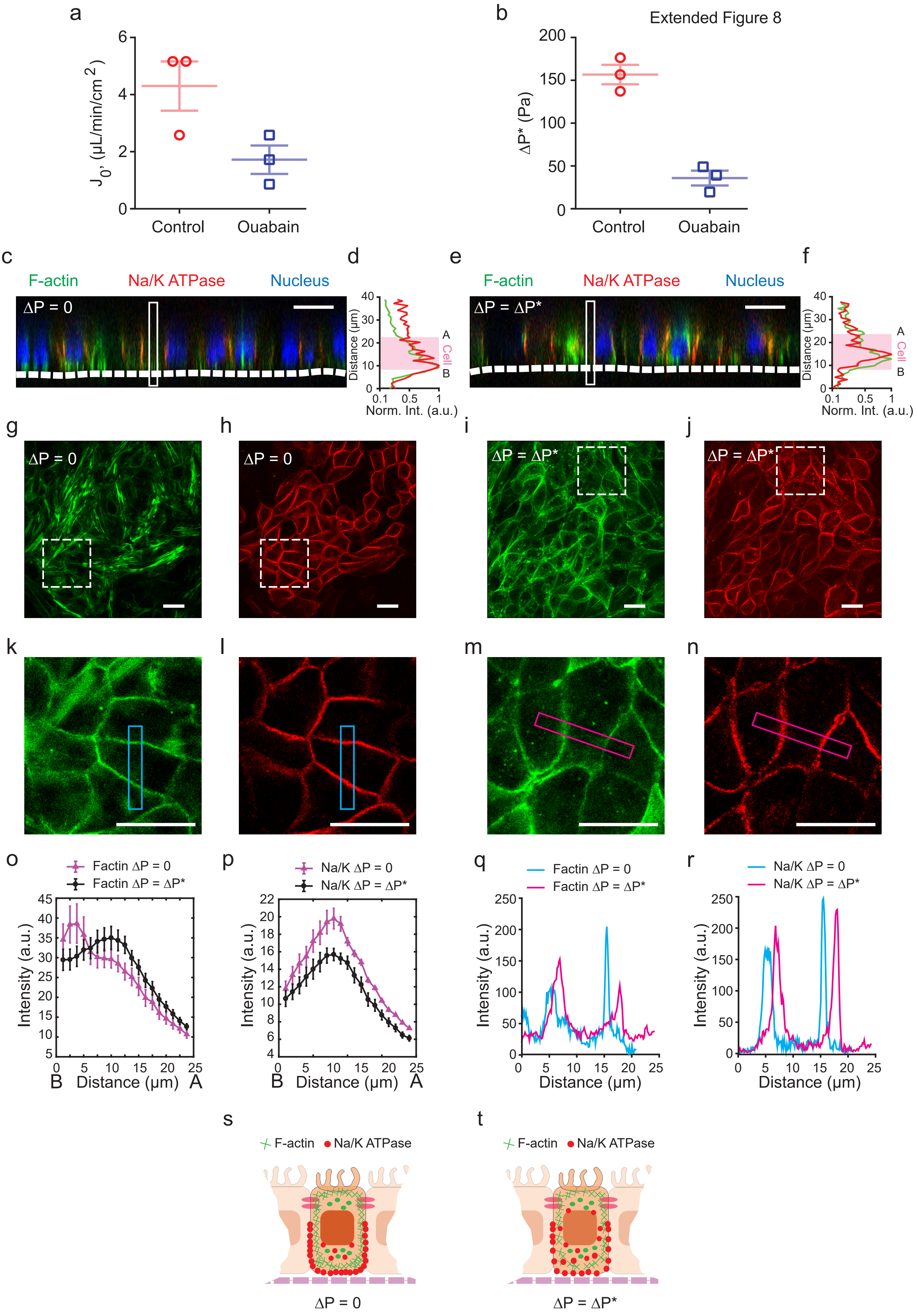

Extended Figure 9

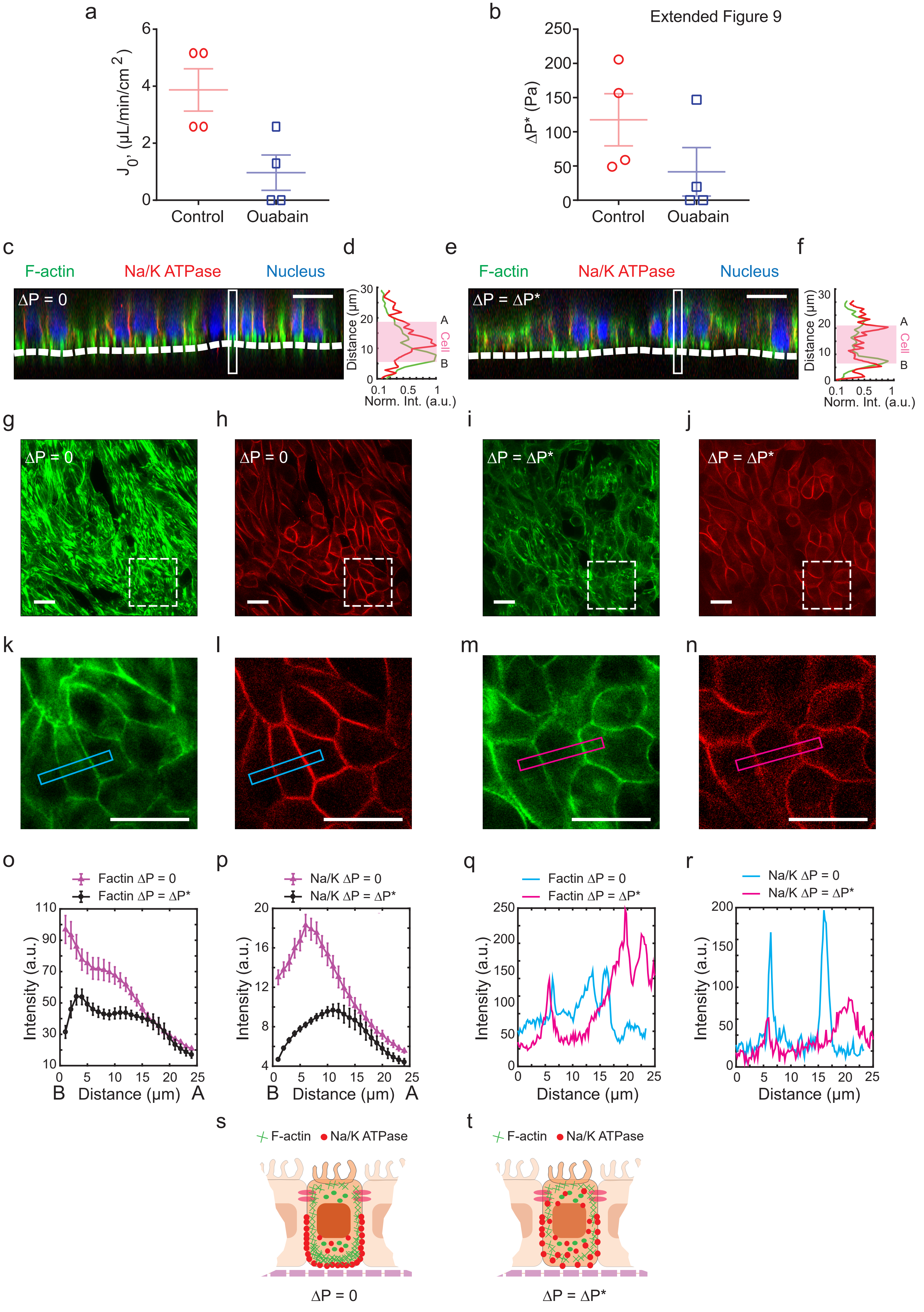

Extended Figure 10

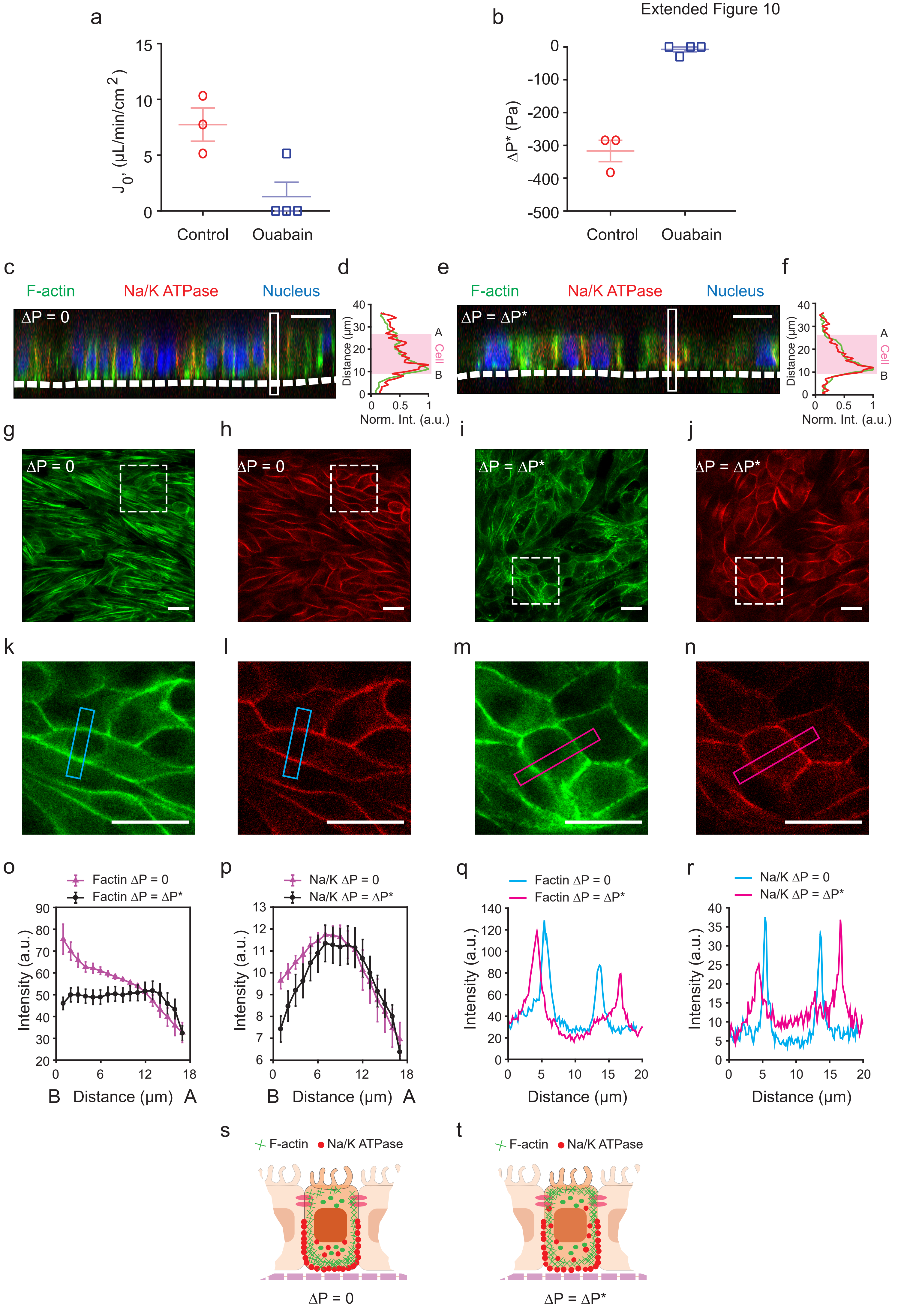
